# Intergenerational transmission of the structure of the auditory cortex and reading skills

**DOI:** 10.1101/2024.09.11.610780

**Authors:** Olga Kepinska, Florence Bouhali, Giulio Degano, Raphael Berthele, Hiroko Tanaka, Fumiko Hoeft, Narly Golestani

## Abstract

High-level cognitive skill development relies on genetic and environmental factors, tied to brain structure and function. Inter-individual variability in language and music skills has been repeatedly associated with the structure of the auditory cortex: the shape, size and asymmetry of the transverse temporal gyrus (TTG) or gyri (TTGs). TTG is highly variable in shape and size, some individuals having one single gyrus (also referred to as Heschl’s gyrus, HG) while others presenting duplications (with a common stem or fully separated) or higher-order multiplications of TTG. Both genetic and environmental influences on children’s cognition, behavior, and brain can to some to degree be traced back to familial and parental factors. In the current study, using a unique MRI dataset of parents and children (135 individuals from 37 families), we ask whether the anatomy of the auditory cortex is related to reading skills, and whether there are intergenerational effects on TTG(s) anatomy. For this, we performed detailed, automatic segmentations of HG and of additional TTG(s), when present, extracting volume, surface area, thickness and shape of the gyri. We tested for relationships between these and reading skill, and assessed their degree of familial similarity and intergenerational transmission effects. We found that volume and area of all identified left TTG(s) combined was positively related to reading scores, both in children and adults. With respect to intergenerational similarities in the structure of the auditory cortex, we identified structural brain similarities for mother-child pairs of the 1st TTG (HG) (in terms of volume, area and thickness for the right HG, and shape for the left HG) and of the lateralization of all TTG(s) surface area for father-child pairs. Both the HG and TTG-lateralization findings were significantly more likely for parent-child dyads than for unrelated adult-child pairs. Furthermore, we established characteristics of parents’ TTG that are related to better reading abilities in children: fathers’ small left HG, and a small ratio of HG to planum temporale. Our results suggest intergenerational transmission of specific structural features of the auditory cortex (not directly linked to children’s reading outcomes); these may arise from genetics and/or from shared environment.

## 1 Introduction

The development of high-level cognitive skills invariably depends on a complex interplay between genetic and environmental factors, and is related to brain structural and functional indices. In the context of brain-behavior relations, inter-individual variability in reading, language, and musical abilities has been repeatedly associated with the structure of the auditory cortex, with prominent accounts proposing that “[t]he neural basis of dyslexia may originate in primary auditory cortex” (Goswami, 2014, p. 3100; in response to Clark et al., 2014). Of particular importance here are the shape, size and asymmetry of the transverse temporal gyrus (TTG) or gyri (TTGs), if there are several (Benner et al., 2017; Turker et al., 2021). The TTG(s) is (are) located within the Sylvian fissure, on the superior surface of the superior temporal gyrus. The most anterior TTG, or the only TTG in the case of a single gyrus, is also known as Heschl’s gyrus (HG) and includes the primary auditory cortex (von Economo and Horn, 1930). Additional TTGs, if present, are part of the planum temporale (PT), which houses the secondary auditory cortex, although the mapping between cytoarchitecture and gross anatomy is not one-to-one. The TTG(s) show(s) high variability in shape and size between hemispheres and individuals: there may be a single gyrus (HG), duplications (with a common stem or fully separated) or multiplications of the TTGs (Geschwind and Levitsky, 1968; Marie et al., 2015). This variability in shape but also in size, has in turn, been frequently associated with individual differences in a range of auditory, language and music-related skills, and with a number of disorders. These include phonetic learning skill (Golestani et al., 2007, 2002) and expertise (Golestani et al., 2011), linguistic pitch learning ability (Wong et al., 2008), musical expertise (Benner et al., 2017) and language aptitude (Ramoser et al., 2025; Turker et al., 2019, 2017). These individual differences can arise from influences both in predisposition and in experience-dependent plasticity, as suggested by earlier work in phonetics experts (Golestani et al., 2011), by our recent work in multilinguals (Kepinska et al., 2025), and in the neurocognitive model of language aptitude proposed by Turker and colleagues (2021). TTG(s) anatomy has also been associated with pathological conditions such as tinnitus (Schneider et al., 2009) and schizophrenia (Takahashi et al., 2022), as well as with (poor) pre-reading and reading skills (Altarelli et al., 2014; Blockmans et al., 2023; Kuhl et al., 2020; Leonard et al., 2001; Serrallach et al., 2016; Sutherland et al., 2012). The latter is the focus of the present study.

Reading skill and disorder have been associated with variations in several structural features of the TTG(s) (gyrification patterns, volume, thickness, surface area, asymmetry, etc.), and as such, there is no clear consensus on which specific anatomical feature is most explanatory of individual differences in reading ability. For example, dyslexia has been associated with differences in gross shape, with more TTG duplications in dyslexic children in the right hemisphere only (Altarelli et al., 2014; Serrallach et al., 2016, with the former study showing these results only in boys), whereas a similar result has previously been found in the left hemisphere, in adults (Leonard et al., 2001). Kuhl and colleagues (2020) showed that a higher local gyrification index around the left HG (reflecting the presence of more involutions and buried cortex) in pre-reading children was predictive of a dyslexia diagnosis three years later. The predictive value of HG surface area and of TTG duplication patterns for later reading was also recently demonstrated by Blockmans and colleagues (2023)). They found that greater surface area of the left HG and of left and right PT, and more left TTG multiplications in pre-reading children were associated with better word reading three years later. With respect to TTG(s) cortical thickness, Clark et al. (2014) found that pre-reading 6-year-olds with thinner HG cortex were more likely to develop dyslexia, while Ma et al. (2015) reported that school-aged children with dyslexia had a thicker right HG and surrounding areas (including superior temporal gyrus and PT). While some inconsistencies remain, these findings overall align with observations of pervasive deficits in the representation and processing of speech sounds in dyslexia. Such deficits are considered by many to be one of the factors at the origin of dyslexia, and the above results show that these can be traced back to auditory cortex neuroanatomical features (Linkersdörferet al., 2012). Indeed, Sutherland et al. (2012) found a higher gray-matter probability in the left HG as a neuroanatomical marker underlying the link between literacy and auditory processing in the temporal domain (again, a relationship that was present in boys but not in girls).

Both genetic and environmental influences on children’s cognition, behavior, and brains can, to some extent, be traced back to familial and parental factors. Van Bergen (2014) suggests that “both parents confer [dyslexia] liability via intertwined genetic and environmental pathways”, and points to intergenerational designs as a promising avenue of research into the etiology of the disorder. Given that TTG morphology has been associated with experiential factors such as bi- and multilingualism (Kepinska et al., 2025; Ressel et al., 2012) but that it has also been shown to have moderate to high heritability (Eyler et al., 2012; Grasby et al., 2020), studying the anatomical features of the TTG(s) across different family members and generations may allow us to better understand the familial transmission (via shared environment and/or of genes) of brain structure and of associated phenotypes. Here, we address this question using a unique brain imaging and behavioral dataset from parents and their children. Understanding the intergenerational, familial transmission of brain structure by examining familial similarity may shed light on the mechanisms of inheritance of complex behavioral traits (Ho et al., 2016), although similarities cannot be pinpointed to genetic inheritance specifically since they could also arise from shared experiential influences. Studies are beginning to unravel the concordance of brain structural indices across family members, including mother-daughter similarities in whole-brain sulcal morphology (Ahtam et al., 2021), corticolimbic circuitry (Dimanova et al., 2023; Yamagata et al., 2016), reading network anatomy (Fehlbaum et al., 2022), and parent-child similarity in whole-brain functional and structural measures (Takagi et al., 2021). Moreover, intergenerational designs have been promising in other research areas, such as socio-communicative skills (Ronconi et al., 2024) and this methodological approach has the potential to overcome the important limitation to study only the children of adults with neurodevelopmental disorders, supporting a more dimensional vision of the neurodiversity of cognitive functions.

Due to the variability in its morphology, the TTG(s) has been notoriously difficult to automatically segment accurately, with template-based analyses largely ignoring individual variability in its shape (Dalboni da Rocha et al., 2020). Recent developments in cortical segmentation of the TTG(s) (Dalboni da Rocha et al., 2023, 2020) provide an anatomically precise, reliable and automatic (i.e., without manual segmentation or subjective decisions) method for delineating the region, and deriving continuously quantified measures describing its shape and size (indexed by volume, surface area and thickness). Each structural measure captures different aspects of cortical anatomy with potentially distinct developmental and functional relevance, with volume giving an overall size estimate, surface area thought to be more strongly genetically influenced and potentially relating to early developmental processes and cortical thickness reflecting maturational changes and being more sensitive to environmental influences and plasticity (Grasby et al., 2020), and finally shape descriptors (e.g., number and configuration of TTGs) capturing qualitative anatomical variation not fully reflected in size-based measures. The present study capitalizes on these recent efforts in cortical segmentation, and includes all of these measures to more comprehensively characterize large inter-individual variability of the auditory cortex anatomy and to better understand which structural features are most closely related to reading skill, and similarity among family members. We aim to determine the degree of familial similarity and intergenerational transmission effects on the anatomy of the auditory cortex in a group of 135 individuals from 37 families, and to identify the precise anatomical features of the TTG(s) that are related to reading ability in children and adults, and across generations (i.e., are there any specific anatomical features of the parents’ TTG(s) that predict children’s reading ability?). Throughout the paper, we refer to *familial similarity* as the resemblance in skills or features of the auditory cortex among biologically related individuals, including parent-child pairs and siblings. This will capture general family-level patterns of similarity that may arise from shared genetics, shared environment, or both. *Intergenerational transmission*, in contrast, will refer specifically to resemblance in traits or brain structures between parent-child dyads. It will focus on whether particular traits are preserved across generations, and thus will exclude sibling-only comparisons. While still correlational, this concept implies a directional relationship from one generation to the next.

## 2 Methods

### 2.1 Participants

MRI data from a total of 135 individuals, including 72 children (*M*_age_ = 8.81, *SD* = 2.11, 35 female), and 63 adults (*M*_age_ = 42.5, *SD* = 5.14, 33 female) were analyzed. In total, there were 37 biological families (65 mother-child, 60 father-child dyads; most families had more than one child), see Table S1 for an overview of the families. Twelve more children were scanned but their data were excluded due to excessive motion and bad segmentation (see Section 2.4 for details). The majority of the participants were right-handed (112:16:3 right:left:ambidextrous ratio); handedness was dummy coded for the analyses, and in 4 cases where the information was missing it was replaced by the mean. Parents’ socioeconomic status (SES) was indexed by the years of education they completed (*M*_mothers_ = 17.08, *SD* = 2.06; *M*_fathers_ = 16.80, *SD* = 2.07); for children, we used an average of the years of completed education of both parents (*M*_SES_ = 16.94, *SD* = 1.74). Across all participating families, 35 individuals reported at least one medical or developmental condition (11 children, 13 mothers, 11 fathers). Among children, the most frequent diagnoses were dyslexia/reading-related difficulties (5 cases) and ADD/ADHD (5 cases), with additional reports of dysgraphia (1), language/working-memory or learning difficulties (2), visual impairment (1), eczema (1), and type I diabetes (1). Among mothers, conditions included dyslexia/reading problems (3), ADD/ADHD (1), depression (2), head injury (1), thyroid cancer (1), ulcerative colitis (1), eosinophilic esophagitis (1), high blood pressure (1), asthma (1), and visual impairment (3). Fathers reported dyslexia/reading difficulties (5), language difficulties (1), ADD/ADHD (1), high blood pressure (2), allergies (1), asthma (1), head injury (1), coronary heart disease (1), and visual impairment (1).

### 2.2 Behavioral data

Reading and reading-related test scores were available for 129 participants (70 children and 59 adults). These included:

1. Sight Word Efficiency (T-SWE) subtest from the Test of Word Reading Efficiency (TOWRE) (Torgesen et al., 1999), in which as many as possible increasingly difficult items from a list of high frequency words including irregular grapheme to phoneme mappings have to be read aloud within 45 seconds;
2. Phonemic Decoding Efficiency (T-PDE) subtest from TOWRE (Torgesen et al., 1999), in which as many as possible increasingly difficult items from a list of high frequency words including pseudo-words have to be read aloud within 45 seconds;
3. Rapid Automatized Naming and Rapid Alternating Stimulus Tests (RAN/RAS) - Letters (RN-LTR) subtest (Wolf and Denckla, 2005), where participants are asked to quickly name fifty individual letters aloud;
4. Woodcock Reading Mastery Tests Revised-Normative Update (WRMT) (Woodcock, 1998): Word Identification (WRMT-WID), consisting of untimed reading of increasingly difficult individual low frequency words;
5. WRMT Word Attack (WRMT-WA) consisting of untimed reading of increasingly difficult individual pseudo-words.

Standard scores^1^ on individual tests were used in the analyses reported in Sections 3.2, 3.3, and 3.5 (see Figure 2).

### 2.3 Data collection procedures

All procedures were approved by the Stanford University Panel on Human Subjects in Medical Research and they were conducted in accordance with its guidelines and regulations. Written informed consent and assent were obtained from parents and children, respectively, after complete description of the study to the participants. Neuroimaging data were collected using a 3T GE-Signa HDxt scanner (GE Healthcare) with a quadrature head coil at the Lucas Center for Imaging at Stanford University. High-resolution T1-weighted anatomical images were acquired with a matrix size of 256*256 and voxel size of 0.86 x 0.86 x 1.2 mm; 156 axial slices; TR = 8.5 ms, TE = 3.4 ms, inversion time = 400 ms; flip angle = 15°; FOV = 22 cm. Images were visually inspected for scanner artefacts and anatomical anomalies.

### 2.4 Segmentation

The T1 images were processed with FreeSurfer’s (version 7.1) structural sub-millimeter pipeline (`recon-all`) (Fischl et al., 2004; Zaretskaya et al., 2018), consisting of motion correction, intensity normalization, skull stripping, and reconstruction of the volume’s voxels into white and pial surfaces. FreeSurfer’s output was visually inspected for segmentation errors, which were caused by excessive motion. We then performed a detailed segmentation of the auditory cortices using an automated toolbox (TASH; Dalboni da Rocha et al., 2020). For morphometric measures, we used `TASH_complete`, which segments and quantifies the cortical structure of all TTG(s) that are identified (Dalboni da Rocha et al., 2023). We performed a visual selection of the gyri segmented by `TASH_complete`, retaining for the analysis only gyri that had a similar orientation as the first TTG (i.e. which we will henceforth refer to as ‘HG’) and excluding gyri along the portion of the superior temporal plane that curved vertically (i.e., within the parietal extension, Honeycutt et al., 2000), when present. From the resulting labels, we exported measures (in native space) of cortical (i.e. grey matter) volume (in mm^3^), surface area (in mm^2^) and mean thickness (in mm). To derive measures of the shape of TTG(s), we used another toolbox, the Multivariate Concavity Amplitude Index (Dalboni da Rocha et al., 2023), which calculates the degree of concavity of each of the gyri segmented by TASH. Following Dalboni da Rocha et al. (2023), lateral concavity values were used in the analysis. We used both an index reflecting the shape of HG alone, and one reflecting the overall shape of all identified TTG(s) (‘lateral multiplication index’, Dalboni da Rocha et al., 2023). The latter was derived by counting all gyri identified by `TASH_complete`, and adding the number of gyri to the lateral concavity values summed across all present TTGs in the respective hemisphere. Higher values for both measures (on HG and on all TTGs) indicate a more complex shape of the TTG(s), e.g., more duplications/multiplications of the TTG (see examples on Figure 1).

**Figure 1.**
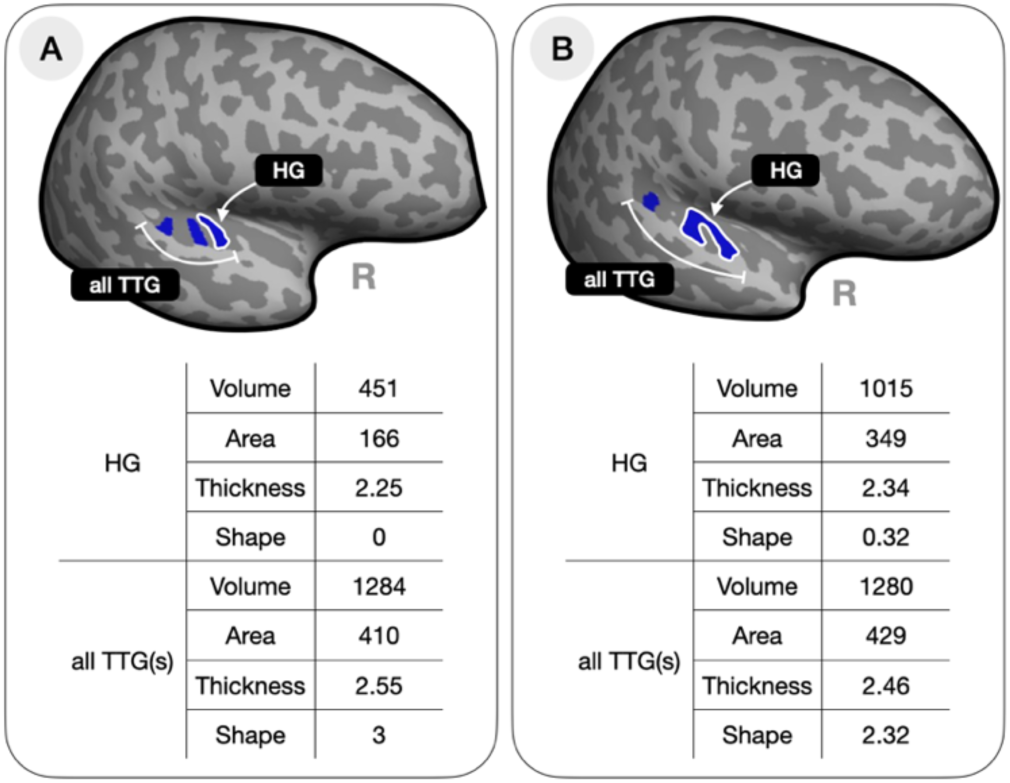
Two examples of the segmented right TTG(s) from the current dataset and the associated anatomical measures used in the statistical analyses: volume (in mm^3^), surface area (in mm^2^), average thickness (in mm) and shape (i.e. the lateral concavity value of the HG as well as the lateral multiplication index for all TTG(s) were extracted using MCAI, (Dalboni da Rocha et al., 2023)). The participant on the left (A) has 3 separate transverse temporal gyri, the one on the right (B) has two gyri, the first being a common stem duplication.

### 2.5 Data analysis

In the statistical analyses, we used two sets of neuroanatomical measures: (1) ones describing only the HG (1^st^ or only TTG) and (2) ones describing all identified TTG(s). Examples of these measures from two participants are shown in Figure 1 below. In addition to the measures of volume, surface area, average thickness and shape of HG and of the TTG(s) per hemisphere, we also computed their lateralization indices (LI). These were calculated using the formula `[left – right]/[left + right]`, with positive values indicating leftward asymmetry. When computing lateralization across all TTG(s), we used the following hemispheric summary measures: (a) for volume and area, we computed the sum of volume and area across all TTG(s) per hemisphere; (b) for thickness, we took the average thickness across the gyri per hemisphere, and (c) for shape, we used the lateral multiplication index, as described above (Section 2.4).

Linear mixed models were used to determine:

(1) the relationships between children’s reading scores and parents’ reading scores (Section 3.2);
(2) the relationships between TTG(s) anatomy and reading scores (Section 3.3);
(3) the familial similarity and intergenerational transmission of TTG anatomy (Section 3.4).

In models testing for intergenerational transmission, likelihood ratio tests were used to compare models that included parental measures with reduced models with covariates of no-interest only. In all analyses, we included fixed factors for covariates of no-interest: age, sex, and SES (indexed by education). Analyses with all brain structural indices as dependent variables included additional fixed factors for participants’ handedness and a quadratic term for age. We accounted for different brain sizes in two ways: total intracranial volume was included as a covariate of no-interest (fixed factor) in analyses where TTG volume, area and thickness were either an independent or dependent variable; in analyses of intergenerational similarity effects (i.e., where children’s anatomical measures were modeled as a functions of measures of their parents), we normalized the anatomical measures for the corresponding whole hemisphere measures (hemispheric volume, surface area and mean hemispheric thickness) so as to be more comparable between parents and children. Specifically, we computed proportional measures for volume, surface area, and cortical thickness in each hemisphere by dividing the regional HG/TTG value by the relevant global hemispheric metric: HG/TTG volume, surface area, and cortical thickness were divided by the total cortical volume, total white matter surface, and mean cortical thickness of the same hemisphere, respectively. This allowed us to account for global anatomical differences between children and adults, which are especially relevant given their differing developmental stages, and ensured that parent-child comparisons reflected regional rather than overall brain morphology.

Familial relationships were coded using a factor variable (“family index”, see Table S1), where all individuals from the same family were assigned the same unique identifier. This variable was included in the analyses to account for the non-independence of observations within families and, in linear mixed models was included as random intercept. In addition, the family index was a critical component of our permutation analyses aimed at testing the specificity of intergenerational similarity effects. To assess whether significant intergenerational similarity effects were specific to true familial relationships rather than driven by spurious correlations in the population, we conducted permutation tests (5,000 iterations). In each permutation, we reassigned parental data (mother and father measures) across families while preserving the original child’s data, including their age, sex, handedness, using the unique family identifier (“family index”). Crucially, parental values were only resampled from individuals belonging to different families (coded by “family index”) to prevent mixing among siblings, and once a parent was selected for a permuted family, they were removed from the sampling pool to avoid reuse. This procedure allowed us to test whether the observed parent-child associations exceeded what would be expected by chance under random familial configurations. All analyses were conducted in R (R Development Core Team, 2015).

In addition, a *random forest* classifier was used to determine:

(4) which, if any, of the parents’ anatomical measures could predict children’s reading (Section 3.5).

Here, we used the `cforest()` function in the `party` package (Hothorn et al., 2006) in R (R Development Core Team, 2015) with 1,000 trees (i.e., `ntree` = 1,000) and four randomly selected predictors considered at each split (i.e., `mtry` = 4). Next, the party’s `varimp()` function (Strobl et al., 2008) was used to compute the conditional permutation importance measures for the predictors, which reflect the impact of each predictor variable on the dependent variable. If the significance (i.e. the *p*-value) of the association between a given predictor and the variable of interest was lower than 0.05, the relevant covariate was included in the predictor’s conditioning scheme. Similarly, we set the `mincriterion` of *p* = .05 to include splits in the calculation of importance.

A summary of all conducted analyses can be found in Section 7.4 of the Supplementary Materials.

## 3 Results

### 3.1 Descriptive statistics

We present the distribution of scores as well as the sample’s means and standard deviations of the collected reading and reading-related test scores in Figure 2 below. Apart from for RN-LTR on which adults scored higher than children (*t*(103.44) = −3.84, *p* = .0002), children’s standard scores were on average higher than those of adults. Significant differences between the groups were noted for T-SWE (*t*(124.97) = −2.55, *p* = .01) and T-PDE (*t*(122.7) = −4.21, *p* < .0001), but not for the WRMT-WA (*t*(116.45) = −1.73, *p* = .09).

**Figure 2.**
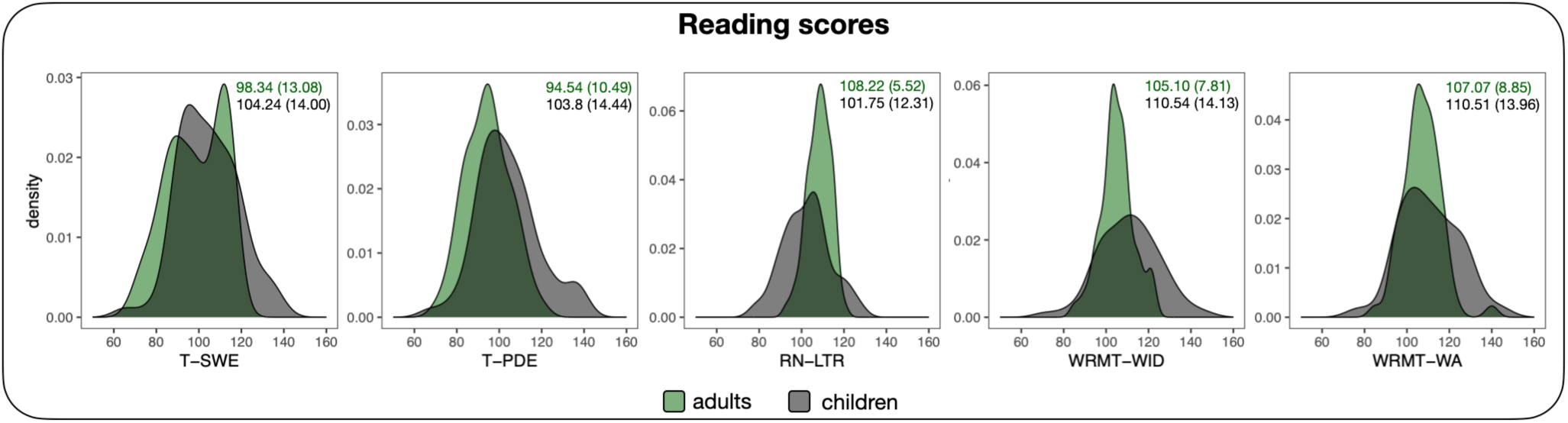
Distributions of standard scores on reading and reading-related tasks for adults (in green) and children (in dark gray): T-SWE = Sight Word Efficiency, T-PDE = Phonemic Decoding Efficiency, RN-LTR = Rapid Automized Naming Letters, WRMT-WID = Word Identification, WRMT-WA = Word Attack. Each panel lists means and standard deviations (in parentheses) of the scores for adults (top rows, in green) and children (bottom rows, in black) separately.

The number of TTG multiplications was highly variable in the sample. In the left hemisphere, we noted only 2 cases of a single gyrus (2 adults); 63 participants had 2 gyri (28 adults, 35 children), 55 had 3 gyri (28 adults, 27 children), and 8 had 4 (1 adult, 7 children). In the right hemisphere, 23 cases of a single gyrus were observed (10 adults, 13 children), 79 participants had 2 gyri (37 adults, 42 children), 24 had 3 gyri (11 adults, 13 children), and 2 had 4 (1 adult, 1 child). Further descriptive data for adults’ and children’s neuroanatomical measures describing the TTG (HG and all TTG(s) separately) can be found in Table 1. Note that all volume, surface and shape measures were significantly left-lateralized (according to a one-sample *t*-test against 0 on the lateralization indices), with the exception of the shape of the HG in adult participants. None of the average thickness values showed significant lateralization.

**Table 1.** Descriptive data for adults’ and children’s measures of HG (HG) and all TTG(s) (volume, area and average thickness) in native space, and their lateralization indices (LI), where positive values indicate leftward lateralization. (*) denotes significantly lateralized measures (according to a one sample t-test against 0). Note that children’s values are consistently higher than adults’, which is in line with normative trajectories of gray matter volume, surface area and cortical thickness reported by e.g., Bethlehem and colleagues (2022).

|  | adults |  |  |  | children |  |  |  |
| --- | --- | --- | --- | --- | --- | --- | --- | --- |
|  | HG |  | all TTG(s) |  | HG |  | all TTG(s) |  |
|  | left | right | left | right | left | right | left | right |
|  | Cortical volume: |  |  |  |  |  |  |  |
| mean | 1227.52 | 990.06 | 2016.92 | 1509.10 | 1413.97 | 1109.65 | 2522.26 | 1738.75 |
| (SD) | (319.65) | (347.39) | (495.65) | (351.55) | (473.54) | (348.69) | (717.74) | (403.37) |
| LI [mean (SD)] | 0.11* (0.18) |  | 0.14* (0.13) |  | 0.11* (0.2) |  | 0.17* (0.14) |  |
|  | Surface area: |  |  |  |  |  |  |  |
| mean | 389.25 | 302.35 | 623.24 | 453.54 | 410.22 | 319.26 | 703.67 | 488.74 |
| (SD) | (108.74) | (95.89) | (158.41) | (99.87) | (133.12) | (89.70) | (175.07) | (100.99) |
| LI [mean (SD)] | 0.12* (0.18) |  | 0.15* (0.13) |  | 0.12* (0.19) |  | 0.17* (0.13) |  |
|  | Average thickness: |  |  |  |  |  |  |  |
| mean | 2.48 | 2.52 | 2.58 | 2.61 | 2.69 | 2.70 | 2.80 | 2.78 |
| (SD) | (0.23) | (0.21) | (0.17) | (0.18) | (0.23) | (0.21) | (0.20) | (0.20) |
| LI [mean (SD)] | -0.009 (0.04) |  | -0.006 (0.03) |  | -0.001 (0.04) |  | 0.004 (0.03) |  |
|  | Shape: |  |  |  |  |  |  |  |
| mean | 0.06 | 0.11 | 2.62 | 2.19 | 0.09 | 0.07 | 2.77 | 2.15 |
| (SD) | (0.07) | (0.13) | (0.61) | (0.59) | (0.10) | (0.11) | (0.70) | (0.66) |
| LI [mean (SD)] | -0.02 (0.68) |  | 0.09* (0.18) |  | 0.24* (0.70) |  | 0.13* (0.21) |  |

### 3.2 Intergenerational transmission of reading skills

We first tested whether children’s performance on the five reading and reading-related tests (T-SWE, T-PDE, RN-LTR, WRMT-WID and WRMT-WA) was related to parents’ performance on the same tests above and beyond children’s demographic variables (age, sex and SES). To establish a relationship between children’s and parents’ reading skills, a series of linear mixed models were fitted to the children’s test scores. To account for sibling relationships (i.e., non-independent observations within the data) family index was included as a random intercept in all models. For each model, a likelihood ratio test was performed to compare models that included parents’ measures with reduced models that did not include them. FDR correction for multiple comparisons was applied to the resulting *p*-values (five from model comparisons including mothers’ data and five with fathers’ data). Figure 3A presents the results of these analyses. Children’s scores were significantly related to mothers’ scores on the same tests for T-PDE (β = 0.63, *t* = 4.76, *p* < .001), WRMT-WID (β = 0.59, *t* = 2.88, *p* = .007) and WRMT-WA (β = 0.36, *t* = 2.34, *p* = .02), and to fathers’ scores on the same tests for T-PDE (β = 0.53, *t* = 2.65, *p* = .01), and WRMT-WID (β = 0.74, *t* = 3.25, *p* = .004). To assess if the above intergenerational similarity effects of reading scores were due to actual familial relationships and not to spurious correlations across our population, we assessed the specificity of these relationships by computing the same models on 5000 permutations of the family index. The analysis confirmed all above significant relations established in the linear mixed models between child’s scores and same test scores for: (1) mother’s T-PDE (*p* = 0.0004), WRMT-WID (*p* = 0.01), WRMT-WA (*p* = 0.04), and (2) father’s T-PDE (*p* = 0.003), and father’s WRMT-WID (*p* = 0.001). In sum, both mother-child and father-child similarities in reading skills were observed, but were more prevalent (i.e., were observed for more of the tested reading measures, and displayed bigger effect sizes, see Figure 3A) for mother-child pairs. Interestingly, they were observed for reading subtests which target phonological decoding skills rather than lexico-semantic, or sight-word reading.

**Figure 3.**
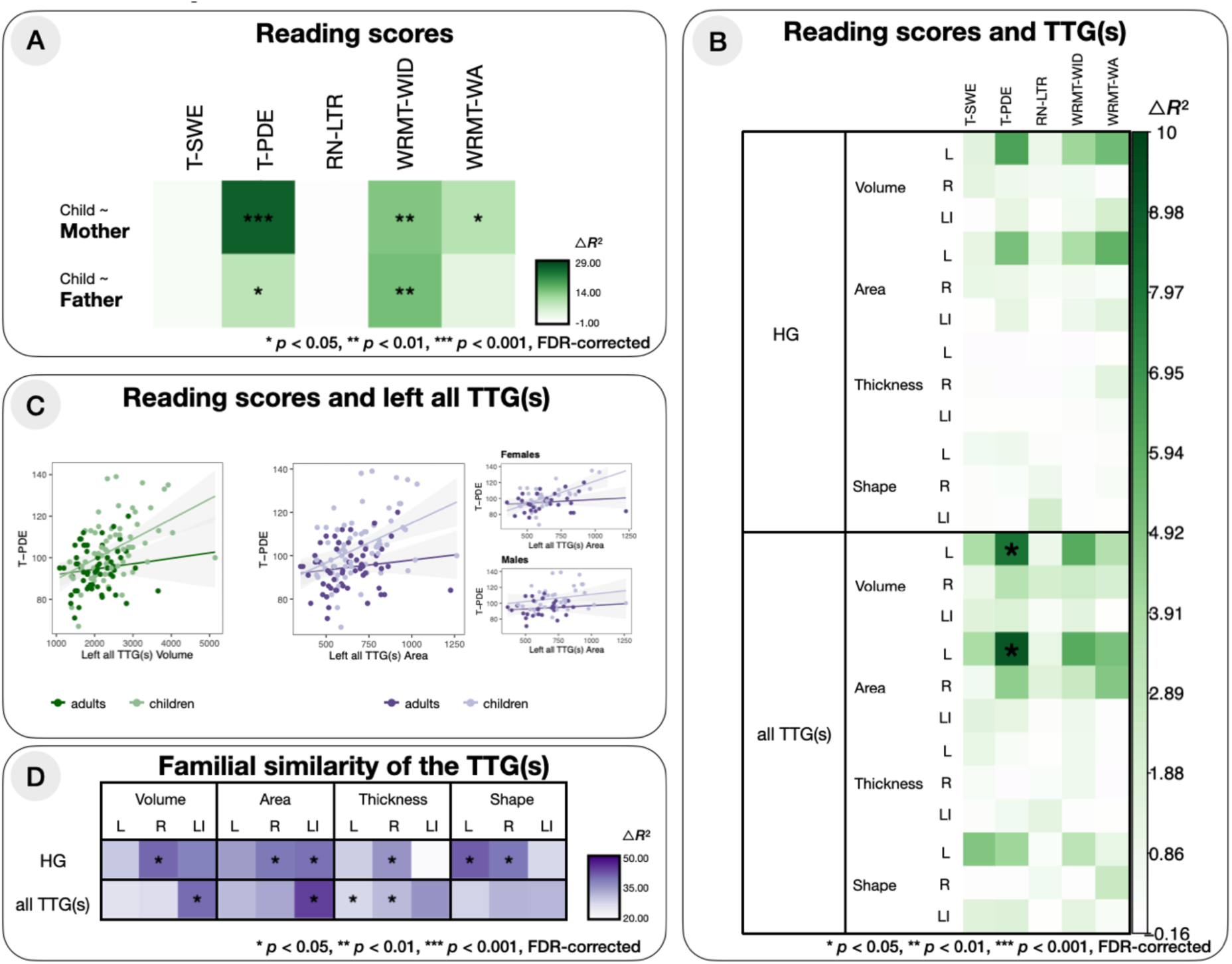
(A) Relationship between parents’ and children’s reading scores on five tests: T-SWE, T-PDE, RN-LTR, WRMT-WID and WRMT-WA. The intensity of the color represents increase in *R*^2^ values between a model with demographic variables only and a model additionally including mothers’ or fathers’ corresponding test scores; *p*-values were obtained from likelihood ratio tests, used to compare the models. (B) Relationship between TTG(s) anatomy and performance on the reading tests across the whole sample, obtained from comparing models with demographic variables to models that additionally included the neuroimaging data; *R*^2^ values and *p*-values were derived and represented similarly as in panel A. (C) Significant relationships between reading tests and neuroanatomical variables. In all plots, darker dots and regression lines refer to adults, and lighter ones refer to children. (D) Familiar similarity of the anatomical measures describing the TTG(s) established by comparing models with demographic variables to models additionally including the family index; *R*^2^ values and *p*-values derived and represented as in panel A.

### 3.3 TTG(s) and reading measures

To explore whether and how the structure of the TTG(s) was related to reading measures in the whole sample, we extracted neuroanatomical measures (volume, surface area, average thickness and shape) of the HG and for all identified TTG(s), from both hemispheres, as well as their lateralization indices. To establish the relationship between the neuroanatomical measures and reading scores, for each reading score, we fitted a linear mixed model predicting this score in a model including a neuroanatomical measure of interest, and a reduced model without it. We compared the two nested models using a likelihood ratio test. In all models, covariates of no-interest (age [linear and quadratic term], sex, SES (indexed by education), handedness and total intracranial volume) were included. FDR correction for multiple comparisons was applied to the resulting *p*-values (120 *p*-values in total: from 24 neuroanatomical measures x 5 reading measures). Significant associations between reading scores and TTG anatomy were observed only in the left hemisphere, for anatomical features in all TTG(s), after correction for multiple comparison (Figure 3B, see also Figure S1 for uncorrected results). Specifically, the models predicting the T-PDE scores by the left all TTG(s) volume (△*R*^2^_Marg._ = 8.15, *p*_FDR_ = .02) and surface area (△*R*^2^_Marg._ = 9.26, *p*_FDR_ = .02), showed a significantly better fit than the models without the TTG(s) indices (after FDR correction). Larger left TTG(s) were associated with better phonemic decoding (volume: β = 0.34, *t* = 2.38, *p* = .001, area: β = 0.34, *t* = 3.37, *p* = .001). Furthermore, in the model predicting T-PDE from left all TTG(s) volume, we observed a significant interaction between left all TTG(s) volume and age (β = −0.24, *t* = −2.26, *p* = .026); the left all TTG(s) area model showed a significant two-way interaction between left all TTG(s) area and age (β = −0.26, *t* = −2.71, *p* = .008), and a significant three-way interaction between left all TTG(s) area, age and sex (β = 0.29, *t* = 2.13, *p* = .036). The significant two-way interactions with age are shown in Figure 3C, where steeper slopes of the relationship between T-PDE and left all TTG(s) measures can be observed for children than for adults. The three-way interaction between left all TTG(s) area, age and sex is driven by the fact that the slope of the relationship between T-PDE and left all TTG(s) area differs between girls, boys, mothers and fathers and is the steepest for girls.

Further, Figure S1 presents the results of the analysis of the relationship between the structure of the TTG(s) and reading measures without FDR correction for multiple comparisons. Here, both the volume of the HG as well as the shape of all TTG(s) show significant relationships with reading measures.

We then ran a follow-up, complementary analysis on the planum temporale (PT). Indeed, additional TTGs by definition belong to the planum, and the anatomy of the PT has been repeatedly linked to reading skills (Altarelli et al., 2014; Blockmans et al., 2023; Serrallach et al., 2016). Therefore, using linear mixed models with age [linear and quadratic term], sex, SES (indexed by education), handedness and total intracranial volume as covariates of no-interest (with family index as a random intercept), we associated the reading measures which were significantly related to ‘all TTG(s)’ volume and area in the previous analysis to the anatomical measures describing the whole PT. We used the PT label as delineated by the Destrieux segmentation (Destrieux et al., 2010), implemented in FreeSurfer. Both the volume (β = 0.16, *t* = 2.07, *p* = .04) of the left PT and its surface area (β = 0.18, *t* = 2.37, *p* = .02) were significantly related to T-PDE. Directly comparing the models predicting the T-PDE scores from ‘all TTG(s)’ volume and area *versus* PT volume and area revealed that ‘all TTG(s)’ measures were better at explaining reading skills, with △*R*^2^_Marg._ of 4.39% (△AIC = −7.78) and 3.55% (△AIC = −6.61) for volume and area respectively. Thus, reading skills may relate more specifically to the anatomy of the gyri encompassed within the HG/PT region of the Sylvian fissure, than to the PT per se. Finally, a two-way interaction between left PT area and sex was observed (β = − 0.32, *t* = −2.36, *p* = .02), with a steeper slope of the relationship between T-PDE and left PT area for females than for males.

### 3.4 Familial similarity in the structure of the TTG(s)

Next, we explored familial similarity and intergenerational transmission of TTG anatomy. A series of linear models were fitted to the participants’ volume, surface area, average thickness and shape of the left and right HG, of all identified left and right TTG(s), and to their lateralization indices. In each of these models, we: (1) modelled the extracted anatomical measures as a function of covariates of no-interest only (participants’ age [linear and quadratic term], sex, handedness, and total intracranial volume), and (2) additionally modelled an index of whether individuals were related or not (i.e., ‘family index’). To establish the familial similarity effect for the four measures, for each we used a likelihood ratio test to compare the model including the family index to the reduced model without it (i.e., with co-variates of no-interest only); FDR correction for multiple comparisons was applied to the resulting *p*-values (24 *p*-values in total). The results of these analyses are summarized in Figure 3D. All models including the family index explained at least 20% more variance than models without (min. △*R*^2^ = 20.44%, max. △*R*^2^ = 43.96%). Including family information significantly improved model fit for the volume (△*R*^2^ = 40.66%, *p*_FDR_ = .024), area (△*R*^2^ = 39.02%, *p*_FDR_ = .044), and thickness (△*R*^2^ = 35.88%, *p*_FDR_ = .024) of the right HG, thickness of all TTG(s) in the left (△*R*^2^ = 27.66%, *p*_FDR_ = .044), and right (△*R*^2^ = 31.40%, *p*_FDR_ = .044) hemispheres, shape of the HG in the left (△*R*^2^ = 41.43%, *p*_FDR_ = .024) and right (△*R*^2^ = 39.11%, *p*_FDR_ = .024) hemispheres, as well as lateralization of the volume (△*R*^2^ = 40.21%, *p*_FDR_ = .044) and surface area of all identified TTG(s) (△*R*^2^ = 43.96%, *p*_FDR_ = .024), and lateralization of the area of the HG (△*R*^2^ = 39.71%, *p*_FDR_ = .050). Familial similarity in TTG anatomy was notably stronger for surface area than thickness, in line with previous reports of higher genetic contributions to shaping surface area (Grasby et al., 2020). Interestingly, across the regions considered, familial similarity was strongest for the right HG, with significant familial contributions in all anatomical features. For all anatomical features of the TTG(s) showing significant familial similarities, we next investigated the intergenerational origin of such similarities by testing parent-of-origin effects.

#### 3.4.1 Intergenerational transmission of the volume, area, and thickness of the 1^st^ right TTG (HG)

To gain further insight into the nature of the familial similarity of the structure of the 1st right TTG, we next examined the relationship between parent and child measures of right HG volume, surface area, and average thickness. For this, we employed linear mixed models, treating the children’s volume, surface and thickness of the right HG as dependent variables, and treating the mothers’ or fathers’ corresponding measures the independent ones, controlling for children’s age [linear and quadratic term], sex, SES, and handedness, and including family index as a random intercept to account for sibling relationships. In total, 6 models were fit (for three different anatomical measures, for the mothers’ and fathers’ data separately). Prior to the analysis, the anatomical measures were normalized for the corresponding whole hemisphere measures (hemispheric volume, surface area and mean hemispheric thickness) so as to be more comparable between parents and children. We found the volume, area, and thickness of children’s 1^st^ right TTG to be significantly related to mothers’ 1^st^ right TTG volume, area, and thickness respectively (volume: β = 0.38, *t* = 2.84, *p* = .009; area: β = 0.30, *t* = 2.33, *p* = .03; thickness: β = 0.38, *t* = 2.87, *p* = .008), as well as children’s thickness to fathers’ thickness values (β = 0.28, *t* = 2.05, *p* = .05), but we did not find relationships between children’s and father’s respective volume and surface area values (volume: β = 0.15, *t* = 1.10, *p* = .28; area: β = 0.16, *t* = 1.14, *p* = .26), see Table 2 and Figure 4.

**Figure 4.**
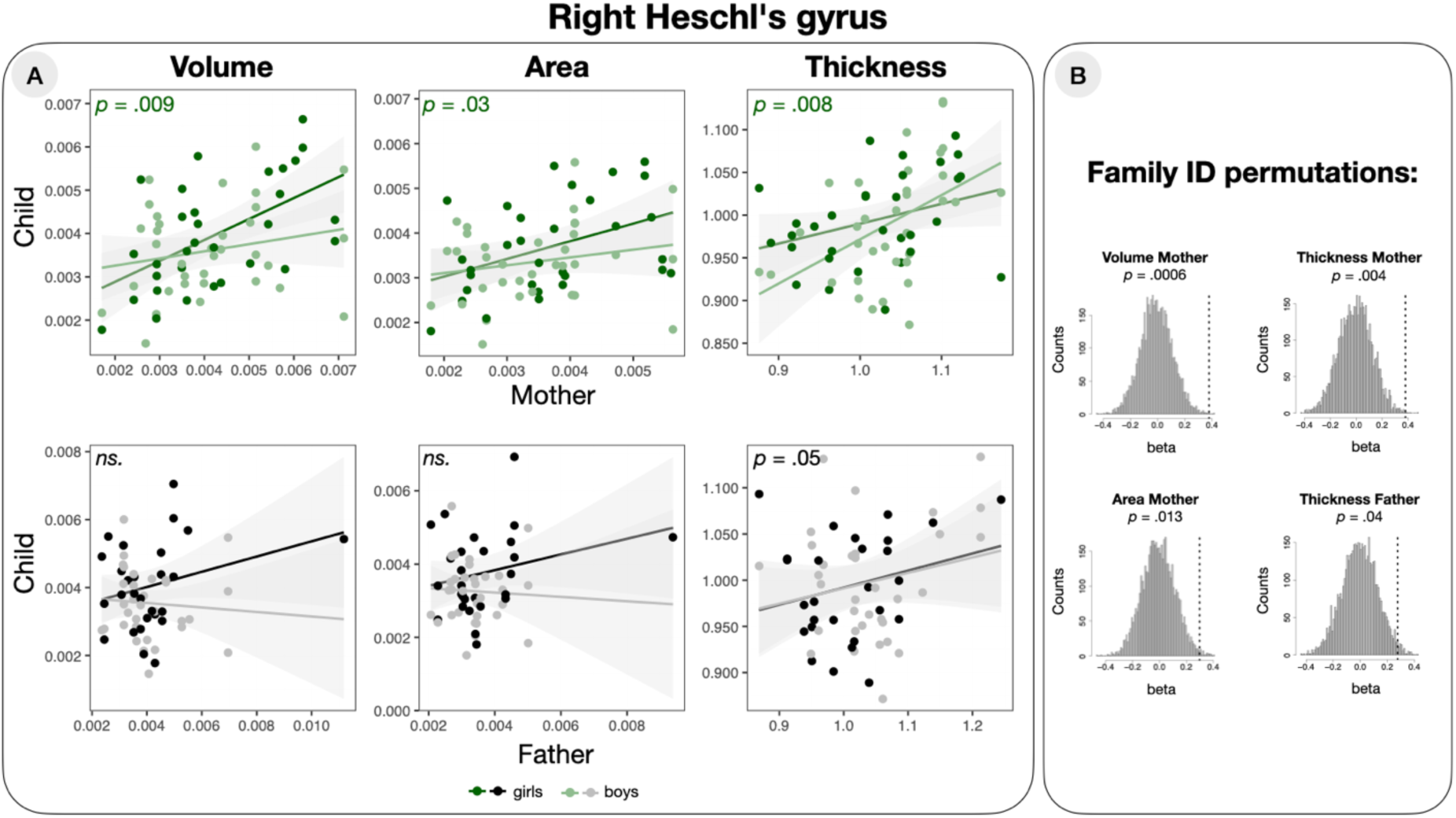
(A) Relationships between right HG volume, surface area and average thickness values of parents and children plotted separately for mothers (top row) and fathers (bottom row); in all plots, darker dots and regression lines refer to girls, lighter dots refer to boys. (B) Distribution of the beta estimates after family permutation. Dotted lines indicate the best estimate of the right HG values of mothers and fathers computed with the correct family labels.

**Table 2.** Linear mixed model parameters for the volume, surface area and average thickness values of children’s right HG as a function of the mothers’ and fathers’ respective HG volume, surface area and average thickness values.

|  |  | Mother |  |  | Father |  |  |
| --- | --- | --- | --- | --- | --- | --- | --- |
|  |  | Volume | Area | Thickness | Volume | Area | Thickness |
| (Intercept) | $\beta$ | 0.12 | 0.12 | 0.16 | 0.22 | 0.24 | 0.19 |
|  | SE | (0.18) | (0.18) | (0.19) | (0.21) | (0.21) | (0.20) |
| Parental Volume/Area/Thickness | $\beta$ | **0.38\*\*** | **0.30\*** | **0.38\*\*** | 0.16 | 0.16 | **0.28\*** |
|  | SE | **(0.13)** | **(0.13)** | **(0.13)** | (0.14) | (0.14) | **(0.13)** |
| Handedness | $\beta$ | -0.10 | -0.12 | 0.01 | -0.12 | -0.10 | -0.25+ |
|  | SE | (0.10) | (0.10) | (0.11) | (0.12) | (0.13) | (0.13) |
| Age | $\beta$ | -0.20+ | -0.19 | -0.20 | -0.16 | -0.14 | -0.23 |
|  | SE | (0.11) | (0.12) | (0.13) | (0.12) | (0.13) | (0.14) |
| Age<sup>2</sup> | $\beta$ | -0.05 | -0.06 | -0.03 | -0.08 | -0.06 | -0.03 |
|  | SE | (0.08) | (0.08) | (0.09) | (0.07) | (0.08) | (0.09) |
| Sex | $\beta$ | -0.25 | -0.26 | -0.25 | -0.38 | -0.47+ | -0.14 |
|  | SE | (0.22) | (0.22) | (0.24) | (0.24) | (0.25) | (0.26) |
| SES (indexed by education) | $\beta$ | -0.03 | -0.02 | -0.15 | -0.17 | -0.18 | -0.09 |
|  | SE | (0.14) | (0.14) | (0.14) | (0.18) | (0.17) | (0.14) |
| *SD (Intercept Family index)* |  | 0.53 | 0.47 | 0.45 | 0.63 | 0.53 | 0.35 |
| *SD (Observations)* |  | 0.70 | 0.75 | 0.83 | 0.71 | 0.80 | 0.86 |
| *Num.Obs.* |  | 65 | 65 | 65 | 60 | 60 | 60 |
| *R<sup>2</sup> Marg.* |  | 0.237 | 0.201 | 0.196 | 0.133 | 0.131 | 0.188 |
| *R<sup>2</sup> Cond.* |  | 0.515 | 0.425 | 0.375 | 0.509 | 0.399 | 0.301 |
| *AIC* |  | 190.6 | 192.2 | 201.6 | 183.0 | 187.5 | 187.0 |
| *BIC* |  | 210.2 | 211.8 | 221.2 | 201.8 | 206.3 | 205.8 |
| *ICC* |  | 0.4 | 0.3 | 0.2 | 0.4 | 0.3 | 0.1 |
| *RMSE* |  | 0.58 | 0.63 | 0.72 | 0.57 | 0.66 | 0.76 |

+ $p < 0.1$ , \* $p < 0.05$ , \*\* $p < 0.01$ , \*\*\* $p < 0.001$

To assess if the above intergenerational similarity effects in the right HG were due to actual familial relationships and not to spurious correlations across our population, we assessed the specificity of these relationships by computing the same models on 5,000 permutations of the family index. The analysis confirmed all significant relations established in the linear mixed models (*p* = 0.0006, *p* = 0.013 and *p* = 0.004, for mother’s volume, area and thickness, and *p* = 0.04 for father’s thickness respectively); see Figure 4B.

To further explore the maternal transmission in the right HG volume and surface area, we checked the potential sex-specificity of this effect, e.g., whether the mother-child similarities were higher for daughters than for sons. Here, we fit further linear mixed models to children’s HG volume and area (with the mothers’ corresponding measures as independent variables, controlling for children’s age [linear and quadratic term], sex, SES (indexed by education), and handedness, and with family index as a random intercept), additionally modelling an interaction between children’s sex and mothers’ HG volume or area. We compared these models with the models without the interaction terms using likelihood ratio tests. These showed that only the interaction model for volume offered a significantly better fit to the data (Χ^2^ = 5.14*, p* = 0.023), and not the one for area (Χ^2^ = 2.16*, p* = 0.14). While it seems that intergenerational similarity of right HG is stronger for mother-daughter pairs, at least for volume, the female specificity of this effect does not seem unequivocal since a visualization of the result for girls and boys separately (Figure 4A) shows similar, positive slopes of the effect for both boys and girls.

#### 3.4.2 Intergenerational transmission of left and right HG shape

To further explain the familial similarity effect of TTG(s) shape reported in Section 3.4, we fit linear mixed models to the right and left HG lateral concavity values as determined by MCAI (Dalboni da Rocha et al., 2023), with mothers’ and fathers’ HG shape indices as independent variables, controlling for children’s age [linear and quadratic term], sex, SES (indexed by education) and handedness, with family index as a random intercept. We found the shape of children’s HG to be significantly related to mothers’ HG shape in the left hemisphere (β = 0.31, *t* = 2.12, *p* = .04), but not for the right hemisphere (β = 0.24, *t* = 1.60, *p* = .12). The relationships between children’s and father’s HG concavity values were not significant (left: β = 0.24, *t* = 1.68 *p* = .11; right: β = 0.17, *t* = 1.20, *p* = .24), see Table 3 and Figure 5.

**Figure 5.**
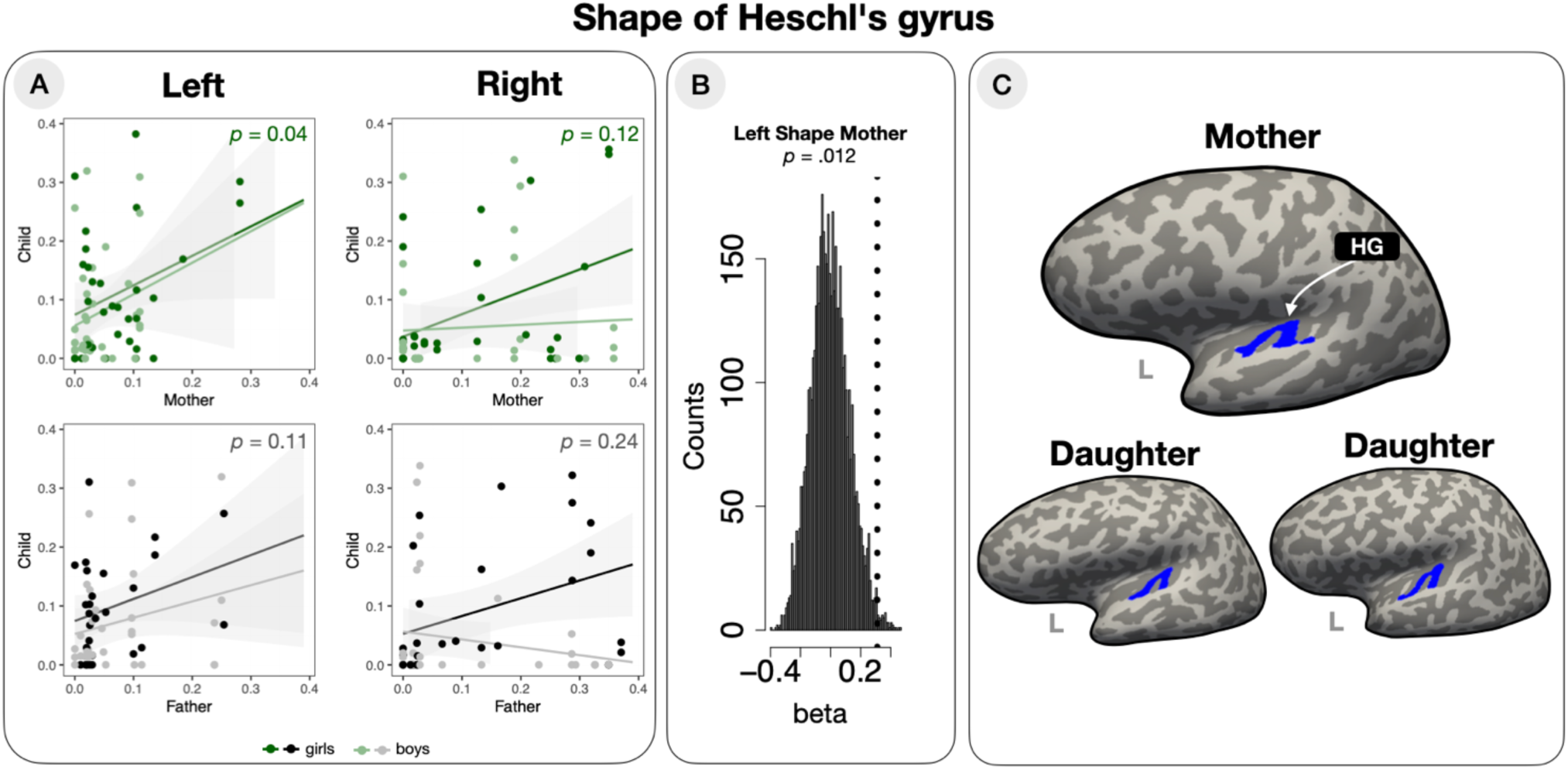
Familial similarity of Heschl’s gyrus shape. (A) Relationship between HG lateral concavity values of parents and children (plotted separately for mothers and fathers) in the left and right hemisphere. In all plots, darker dots and regression lines refer to girls, lighter dots refer to boys. (B) Distribution of the beta estimates after family permutation. Dotted line indicates the best estimate of the HG concavity of mothers computed with the correct family labels. (C) An example of left HG shape similarity between family members.

**Table 3.** Multiple regression model parameters for the concavity values of children’s HG as a function of mothers’ and fathers’ HG concavity values.

|  |  | Mother |  | Father |  |
| --- | --- | --- | --- | --- | --- |
|  |  | Left | Right | Left | Right |
| (Intercept) | $\beta$ | 0.01 | 0.11 | -0.02 | 0.18 |
|  | SE | (0.21) | (0.20) | (0.20) | (0.20) |
| Parent concavity | $\beta$ | **0.31\*** | 0.24 | 0.24+ | 0.18 |
|  | SE | **(0.15)** | (0.15) | (0.14) | (0.15) |
| Handedness | $\beta$ | -0.14 | 0.10 | -0.03 | -0.02 |
|  | SE | (0.12) | (0.10) | (0.13) | (0.12) |
| Age | $\beta$ | -0.06 | -0.01 | -0.06 | 0.06 |
|  | SE | (0.14) | (0.11) | (0.13) | (0.13) |
| Age<sup>2</sup> | $\beta$ | 0.13 | -0.19* | 0.10 | -0.09 |
|  | SE | (0.09) | (0.07) | (0.08) | (0.08) |
| Sex | $\beta$ | -0.20 | -0.09 | -0.31 | -0.35 |
|  | SE | (0.26) | (0.22) | (0.24) | (0.24) |
| SES (indexed by education) | $\beta$ | 0.04 | -0.03 | -0.01 | -0.22 |
|  | SE | (0.15) | (0.16) | (0.15) | (0.17) |
| *SD (Intercept Family index)* |  | 0.42 | 0.70 | 0.48 | 0.52 |
| *SD (Observations)* |  | 0.90 | 0.66 | 0.77 | 0.75 |
| *Num.Obs.* |  | 65 | 65 | 60 | 60 |
| *R<sup>2</sup> Marg.* |  | 0.143 | 0.133 | 0.108 | 0.098 |
| *R<sup>2</sup> Cond.* |  | 0.294 | 0.597 | 0.356 | 0.393 |
| *AIC* |  | 208.0 | 194.3 | 181.9 | 181.8 |
| *BIC* |  | 227.6 | 213.9 | 200.7 | 200.7 |
| *ICC* |  | 0.2 | 0.5 | 0.3 | 0.3 |
| *RMSE* |  | 0.79 | 0.51 | 0.65 | 0.62 |

+ $p < 0.1$ , \* $p < 0.05$ , \*\* $p < 0.01$ , \*\*\* $p < 0.001$

Similarly to the analysis of HG volume, area and average thickness values, we computed 5,000 permutations of the family index to assess the specificity of the familial relationship of left HG shape. The analysis confirmed that the concavity of children’s HG was significantly related to their mothers’ HG concavity in the left hemisphere (*p* = 0.014), see Figure 5B.

Last, we explored the female-specificity of HG similarity by fitting a further linear mixed model to children’s HG shape values (with the mothers’ corresponding measure as independent variable, controlling for children’s age [linear and quadratic term], sex, SES (indexed by parental education), and handedness, and with family index as a random intercept), with additionally modelling an interaction between children’s sex and mothers’ HG shape values. We compared this model with the model without the interaction term using a likelihood ratio test, which showed that the interaction model did not offer a significantly better fit to the data (Χ^2^ = 0.37*, p* = 0.54). Therefore, the left HG shape intergenerational transmission did not appear sex-specific.

#### 3.4.3 Intergenerational transmission of TTG lateralization

We further explored the familial similarity of the lateralization of all TTG(s) (see Figure 6A) by fitting further linear models to the children’s lateralization indices of TTG(s) volume and surface area, with mothers’ or fathers’ corresponding lateralization values as independent variables, controlling for children’s age [linear and quadratic term], sex, SES (indexed by parental education), and handedness, see Table 4. We found the degree of lateralization of the surface area of children’s all TTG(s) to be significantly and positively related to the lateralization of the fathers’ all TTG(s) surface area (β = 0.42, *t* = 3.00, *p* = .005), but not to that of the mothers (β = 0.04, *t* = 0.26, *p* = .80), see Figure 6B. We found no significant effects in the follow-up analyses on lateralization of volume (mother: β = 0.03, *t* = 0.20, *p* = .84; father: β = 0.26, *t* = 1.74, *p* = .09). Permutation analyses confirmed the specificity of this effect to parent-child pairs: the lateralization of the surface area of all TTG(s) of the fathers was significantly related to that of their children (*p* = 0.0006), see Figure 6C. Nested linear mixed models comparison with a likelihood ratio test revealed that a model including an interaction term between fathers’ lateralization index of all TTG(s) area and children’s sex (in addition to demographic variables (age [linear and quadratic term], sex, SES and handedness), random intercept for family index and fathers’ lateralization index of all TTG(s) area) did not have a better fit to children’s all TTG(s) area lateralization (Χ^2^ = 0.25*, p* = 0.61). The male-specificity of familial similarity of the lateralization of all TTG(s) area could therefore not be confirmed.

**Figure 6.**
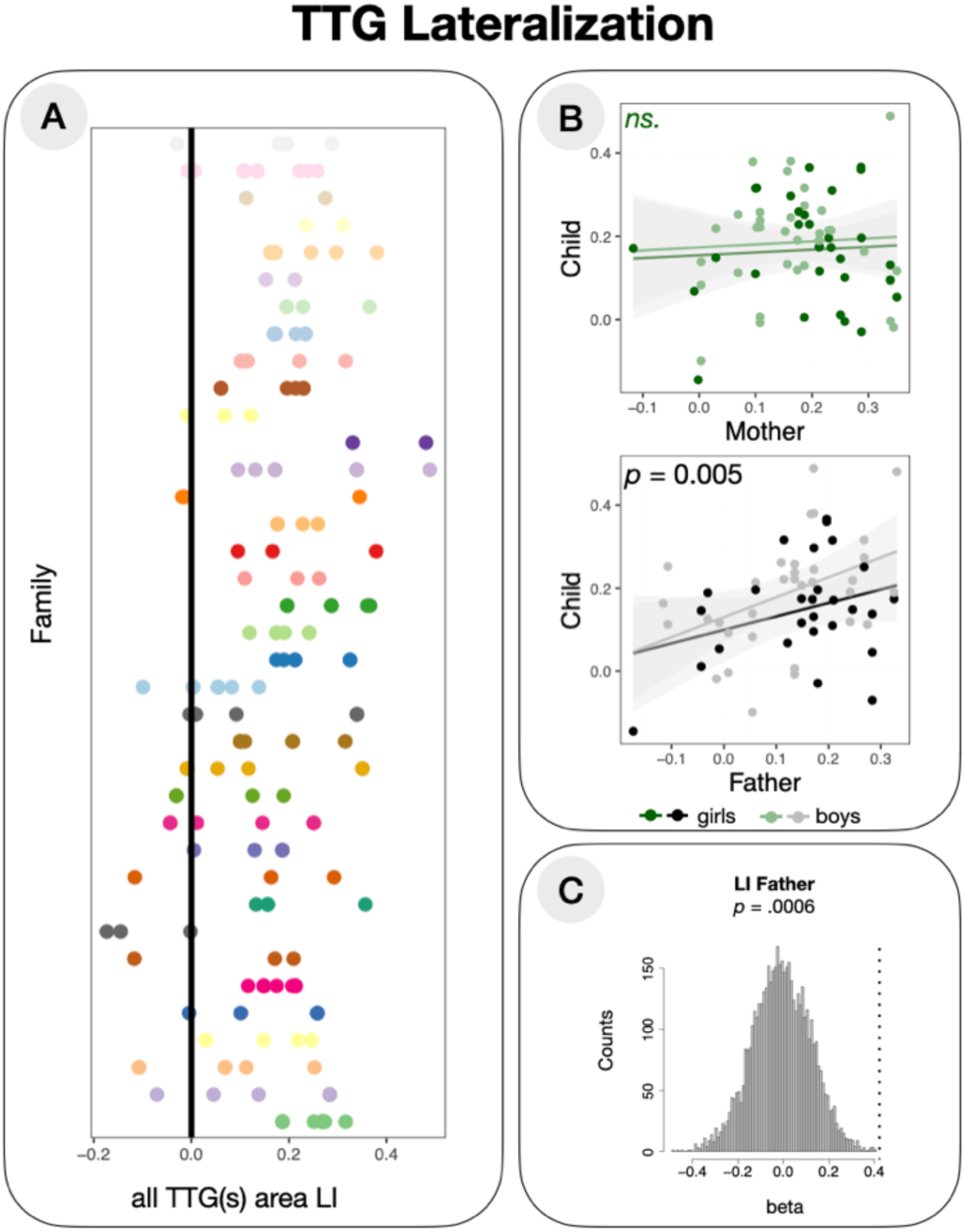
Familial similarity of TTG(s) lateralization. (A) Lateralization indices of TTG(s) (*x*-axis) for 37 different families (*y*-axis); each family is plotted in a different color. (B) Relationship between lateralization index of TTG(s) surface area of parents and children, plotted separately for mothers and fathers. In both plots, darker dots and regression lines refer to girls, lighter dots refer to boys. (C) Distribution of the beta estimates after family permutation. Dotted line indicates the best estimate of lateralization of TTG(s) area of fathers computed with the correct family labels.

**Table 4.** Multiple regression model parameters for the lateralization indices of children’s surface area of all TTG(s) as a function of mothers’ and fathers’ surface area of all TTG(s) lateralization indices.

|  |  | Mother |  | Father |  |
| --- | --- | --- | --- | --- | --- |
|  |  | LI Volume | LI Area | LI Volume | LI Area |
| (Intercept) | $\beta$ | -0.06 | -0.17 | -0.17 | -0.27 |
|  | SE | (0.20) | (0.20) | (0.22) | (0.21) |
| Parent LI | $\beta$ | 0.03 | 0.04 | 0.26+ | **0.42\*\*** |
|  | SE | (0.14) | (0.14) | (0.15) | **(0.14)** |
| Handedness | $\beta$ | 0.07 | 0.04 | -0.02 | -0.04 |
|  | SE | (0.11) | (0.11) | (0.13) | (0.13) |
| Age | $\beta$ | -0.28* | -0.31* | -0.21 | -0.21 |
|  | SE | (0.13) | (0.12) | (0.14) | (0.13) |
| Age<sup>2</sup> | $\beta$ | 0.03 | 0.08 | 0.00 | 0.04 |
|  | SE | (0.08) | (0.08) | (0.08) | (0.08) |
| Sex | $\beta$ | 0.16 | 0.24 | 0.30 | 0.39 |
|  | SE | (0.24) | (0.24) | (0.26) | (0.25) |
| SES (indexed by education) | $\beta$ | -0.13 | -0.12 | 0.01 | 0.05 |
|  | SE | (0.16) | (0.16) | (0.17) | (0.16) |
| SD (Intercept Family index) |  | 0.62 | 0.61 | 0.62 | 0.58 |
| SD (Observations) |  | 0.77 | 0.78 | 0.76 | 0.81 |
| Num.Obs. |  | 65 | 65 | 65 | 60 |
| R<sup>2</sup> Marg. |  | 0.074 | 0.086 | 0.106 | 0.113 |
| R<sup>2</sup> Cond. |  | 0.439 | 0.431 | 0.459 | 0.417 |
| AIC |  | 195.5 | 203.7 | 202.0 | 190.9 |
| BIC |  | 210.8 | 223.3 | 221.5 | 209.7 |
| ICC |  | 0.4 | 0.4 | 0.4 | 0.3 |
| RMSE |  | 0.63 | 0.64 | 0.62 | 0.66 |

+ $p < 0.1$ , \* $p < 0.05$ , \*\* $p < 0.01$ , \*\*\* $p < 0.001$

#### 3.4.4 Lack of intergenerational transmission of other features showing familial similarity

We also performed follow-up analyses on the remaining results that showed significant familial similarity effects in the first exploratory analysis reported in Section 3.4. These included the thickness of the left and right all TTG(s), and the lateralization of the area of the HG. Here, again, linear mixed models were used to relate these measures in children to those of their mothers and fathers. With respect to the measure of average thickness of all left TTG(s), neither the mothers’ (β = 0.25, *t* = 1.78, *p* = .09) nor the fathers’ (β = 0.24, *t* = 1.72, *p* = .10) measures were significantly related to those of the children. This was also the case for all right TTG(s) (mothers: β = 0.19, *t* = 1.77, *p* = .08; fathers: β = 0.28, *t* = 1.98, *p* = .053). Similarly, neither the mothers’ (β = 0.22, *t* = 1.53, *p* = .14) nor the fathers’ (β = 0.21, *t* = 1.47, *p* = .15) lateralization of the area of the HG were significantly related to the children’s lateralization of the area of the HG. Therefore, the effect of “family” for the above neuroanatomical measures reported in Section 3.4 might have been driven by anatomical similarities between the siblings, rather than between parents and children.

### 3.5 Intergenerational effects on reading ability: is children’s reading related to parents’ neuroanatomy?

As a last step in our analyses, we aimed to determine whether mothers’ or fathers’ neuroanatomical measures could predict children’s reading. Given the large number of neuroanatomical variables describing the TTG(s), and given that several different brain measures were either related to reading performance (Section 3.3) or showed neuroanatomical concordance between parents and children (Section 3.4), we opted for an exploratory approach using a *random forests* classifier (Breiman, 2001). Here, we used children’s T-PDE scores as a dependent variable, given their association with TTG(s) anatomy established in Section 3.3. We grew two random forests modelling children’s T-PDE in terms of all available (1) mothers’ and (2) fathers’ anatomical measures describing the HG and all TTG(s). Prior to the analysis, the anatomical measures were normalized for the corresponding whole hemisphere measures (hemispheric volume, surface area and mean hemispheric thickness). Conditional permutation importance scores were computed for all models, and are presented in Figure 7A. The dotted vertical lines demarcate the permutation scores that can differ from zero due to randomness alone; variables having importance scores to the left of this line can be considered not to be relevant predictors.

**Figure 7.**
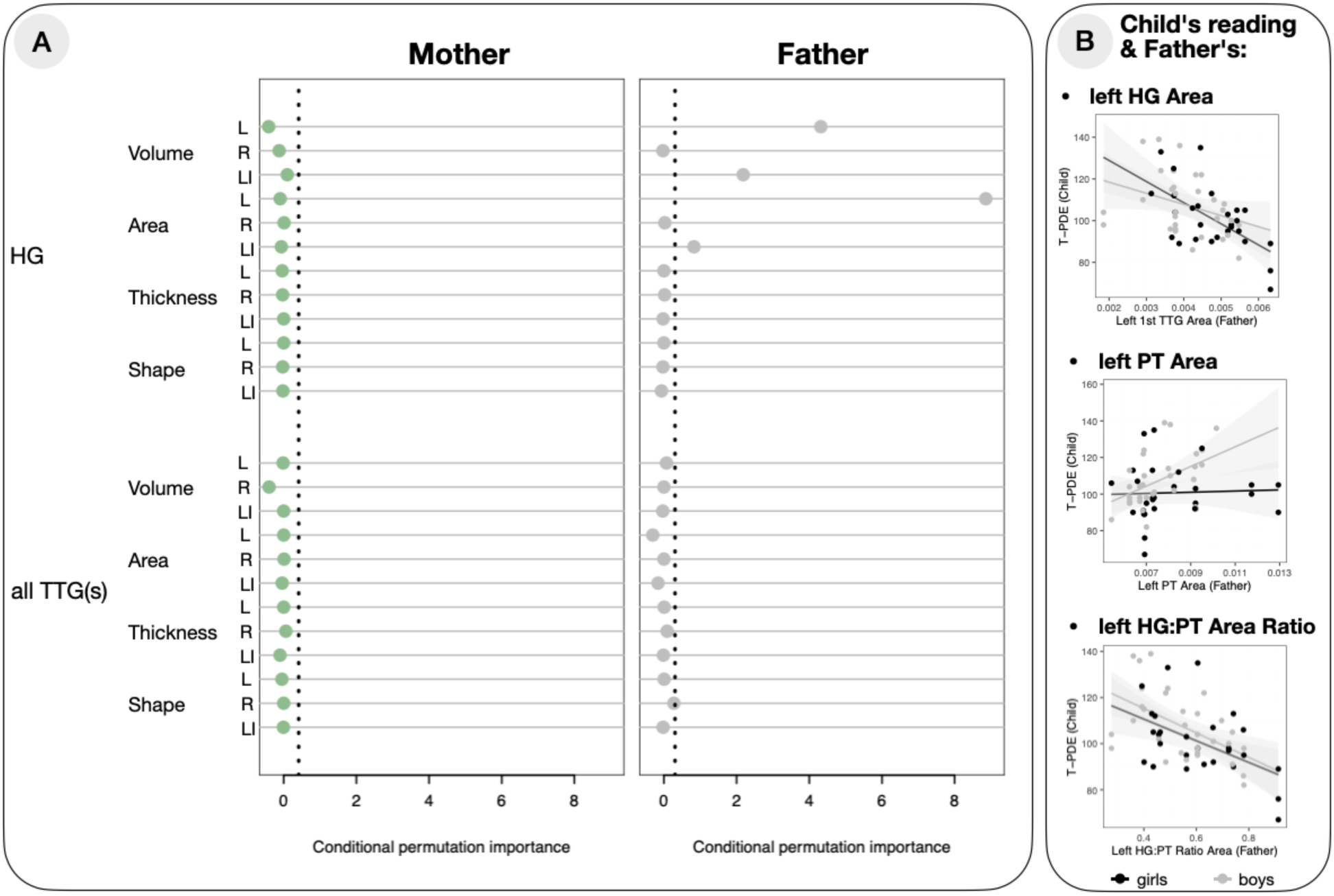
Results of the random forests analysis. (A) Conditional permutation importance for mothers’ and fathers’ neuroanatomical TTG measures in predicting children’s reading skills. (B) Children’s reading skills as a function of fathers’ anatomical variables based on the follow-up analyses (additional plots for father’s volume variables can be found in Supplementary Materials, Figure S2).

The model with mothers’ variables did not point to any features of their TTG(s) to be relevant predictors of children’s reading ability, but several variables in the model with fathers’ data did. The variables with the highest conditional permutation importance in case of fathers were volume and area of the left HG, as well as HG lateralization indices of volume and area; all had conditional permutation importance significantly different from zero, see Figure 7A. To validate these effects and to determine their direction, we fitted separate linear mixed models to children’s T-PDE scores, each with one of the relevant predictors as independent variables (controlling for children’s sex, age and SES, indexed by parental education), and with family index as random intercept. Fathers’ left volume and area were both negatively related to children’s reading (volume: β = −0.40, *t* = −2.91, *p* = .008; area: β = −0.46, *t* = −3.45, *p* = .002), indicating that the smaller the father’s left HG, the better the child’s reading scores. Fathers’ lateralization of HG (both for volume and area) was not significantly related to the children’s reading (volume: β = −0.29, *t* = −1.27, *p* = .21; area: β = −0.20, *t* = −1.32, *p* = .19), see Figure7B and Table 5.

**Table 5.** Multiple regression model parameters for children’s reading ability as a function of the mothers’ and fathers’ anatomical variables having shown the highest conditional permutation importance in the random forests analysis.

|  |  | Father<br>left HG<br>Volume | Father<br>left HG Area | Father<br>LI HG<br>Volume | Father<br>LI HG Area |
| --- | --- | --- | --- | --- | --- |
| (Intercept) | $\beta$ | -0.06 | -0.01 | -0.07 | -0.07 |
|  | SE | (0.19) | (0.19) | (0.21) | (0.21) |
| Father's measure | $\beta$ | **-0.40\*\*** | **-0.46\*\*** | -0.19 | -0.20 |
|  | SE | **(0.14)** | **(0.13)** | (0.15) | (0.15) |
| Child's age | $\beta$ | -0.20+ | -0.20+ | -0.19 | -0.19 |
|  | SE | (0.12) | (0.12) | (0.12) | (0.12) |
| Child's sex | $\beta$ | 0.25 | 0.15 | 0.30 | 0.31 |
|  | SE | (0.26) | (0.26) | (0.27) | (0.27) |
| SES (indexed by education) | $\beta$ | 0.15 | 0.16 | 0.12 | 0.12 |
|  | SE | (0.14) | (0.14) | (0.17) | (0.17) |
| *SD (Intercept Family index)* |  | 0.37 | 0.31 | 0.56 | 0.56 |
| *SD (Observations)* |  | 0.84 | 0.84 | 0.82 | 0.82 |
| *Num.Obs.* |  | 58 | 58 | 58 | 58 |
| *R<sup>2</sup> Marg.* |  | 0.259 | 0.303 | 0.130 | 0.134 |
| *R<sup>2</sup> Cond.* |  | 0.382 | 0.385 | 0.409 | 0.411 |
| *AIC* |  | 172.8 | 170.3 | 178.4 | 178.2 |
| *BIC* |  | 187.2 | 184.8 | 192.8 | 192.6 |
| *ICC* |  | 0.2 | 0.1 | 0.3 | 0.3 |
| *RMSE* |  | 0.75 | 0.76 | 0.69 | 0.69 |

+ $p < 0.1$ , \* $p < 0.05$ , \*\* $p < 0.01$ , \*\*\* $p < 0.001$

To explore sex specificity of these effects, we performed a series of nested model comparisons with likelihood ratio tests for models including (1) left HG Volume, (2) left HG Area, (3) LI HG Volume, or (4) LI HG Area, additionally including an interaction term between fathers’ measures and child’s sex. The results did not point to sex specificity of any of the effects (left HG Volume: Χ^2^ = 2.25*, p* = 0.13, left HG Area: Χ^2^ = 0.88*, p* = 0.35, LI HG Volume: Χ^2^ = 0.17*, p* = 0.68, LI HG Area: Χ^2^ = 0.60*, p* = 0.44) Furthermore, we again tested for familial specificity of the effect of father’s left HG volume and area on children’s reading with permutation analyses. These revealed that the relationship between fathers’ anatomical measures (volume and surface area of the HG) was significantly more related to their own children’s reading scores than to non-related children’s reading scores (*p* = 0.002 and *p* = 0.0006 for fathers’ volume and surface area, respectively).

Given the surprising direction of the intergenerational effects on children’s reading ability, we set out to further understand why fathers’ small left HG would be associated with better reading skills in children. One possibility is that fathers’ small left HG goes hand in hand with more additional TTGs, or with a larger PT (region exclusively or mostly including non-primary auditory areas). Using linear models, we therefore tested (1) whether a smaller left HG in fathers was associated with a greater likelihood of having additional TTGs (reflected by a higher lateral multiplication index for left all TTG(s)), and (2) if there was a negative relationship between the volume and/or area of fathers’ left HG and those of their PT. The lateral multiplication index for left all TTG(s) was not significantly associated with either the volume (β = 0.08, *t* = 0.40, *p* = .69), or the surface area (β = −0.05, *t* = 0.19, *p* = .78) of the left HG in fathers. Surface area of fathers’ left HG showed, however, a positive relationship with the surface area of their PT (β = 0.18, *t* = 2.036, *p* = .05), which was just above the conventional significance threshold, but volume of HG was not related to PT volume (β = 0.20, *t* = 1.08, *p* = .29).

We therefore further evaluated the relationships between children’s reading and the characteristics of their fathers’ auditory cortex regions using linear mixed models (with children’s age, sex and SES, indexed by parental education) as covariates of no-interest and family index as a random intercept). Here, in separate models, we included both Heschl’s gyrus and planum temporale as independent variables, as well as how much space both structures occupy relatively to each other (i.e., HG:PT ratio) (see Serrallach et al., 2016 for findings of a decreased left HG:PT ratio in dyslexia). We concentrate on the surface area here, but see Supplementary Materials for convergent results for the volume. First, while fathers’ PT area alone did not significantly predict children’s T-PDE scores (β = 0.11, *t* = 0.64, *p* = .53), fathers’ HG:PT ratio did indeed significantly and strongly predict children’s reading (β = −0.48, *t* = −3.57, *p* = .002), negatively. The HG:PT ratio effect was not sex-specific: a nested model comparison with a likelihood ratio test did not show a better fit for a model additionally including an interaction term between fathers’ HG:PT ratio and child’s sex (Χ^2^ = 0.44*, p* = 0.51). Conversely, however, we noted a significantly better fit for a model with an interaction term between fathers’ PT area and child’s sex over a model with fathers’ PT area only (Χ^2^ = 5.34*, p* = 0.02): while fathers’ PT area did not significantly predict children’s T-PDE overall (as reported above), and was not significantly related to girls’ reading (β = −0.12, *t* = −0.65, *p* = .53), it was so for boys only (β = 0.66, *t* = 2.40, *p* = .03).

Permutation analyses confirmed familial specificity of the effect of fathers’ HG:PT ratio on children’s reading (*p* < .0001), and fathers’ PT area on boys’ reading (*p* = .005). Directly comparing the models predicting children’s T-PDE with either father’s left HG surface area, or fathers’ HG:PT ratio as independent variables showed that the model with HG:PT ratio offered a better fit to the data with △*R*^2^_Marg._ of 0.9% (△AIC = −0.81). For boys, the model with the lowest AIC was still the HG:PT ratio model (94.70 *versus* 98.99 for father’s left HG surface area, and 96.77 for fathers’ left PT), despite the significant effect of fathers’ left PT surface area on boys’ reading.

In addition, to confirm father-specific transmission, we ran the same three models but with mothers’ HG, PT and HG:PT ratio surface area instead of fathers’. We found no significant relationships between mothers’ neuroanatomical measures and children’s reading (HG: β = 0.11, *t* = 0.78, *p* = .44; PT: β = 0.13, *t* = 0.92, *p* = .37; HG:PT ratio: β = 0.02, *t* = 0.12, *p* = .91). Associating children’s own neuroanatomical measures (surface area/volume of left HG, left PT and left HG:PT ratio) with their reading (controlling for age, sex, SES (indexed by parental education), handedness and total intracranial volume, and with family index as a random intercept) in linear mixed models pointed only to the left HG as a significant predictor of reading (area: β = 0.26, *t* = 2.28, *p* = .03; volume: β = 0.31, *t* = 2.60, *p* = .01). Neither left PT (area: β = 0.16, *t* = 1.43, *p* = .16; volume: β = 0.10, *t* = 0.88, *p* = .39), nor HG:PT ratio (area: β = 0.09, *t* = 0.68, *p* = .50; volume: β = 0.19, *t* = 1.44, *p* = .16) were significantly associated with children’s reading. Lastly, of note, there was no significant relationship between the father’s left HG, PT, or HG:PT ratio and their own T-PDE scores (left HG: β = −0.32, *t* = −1.64, *p* = .11; left PT: β = −0.08, *t* = −0.36, *p* = .72; left HG:PT: β = −0.30, *t* = −1.33, *p* = .19), according to linear models controlling for age, handedness, SES (indexed by education) (and total intracranial volume for left HG and PT models).

In sum, we observed strong intergenerational, father-specific effects on offspring’s reading, manifested by the relative sizes of left primary (HG) and secondary (PT) auditory areas: a small left HG:PT ratio in fathers is related to worse reading in children. Moreover, fathers’ PT area was positively associated with reading in boys only.

## 4 Discussion

The purpose of the present study was to explore intergenerational similarity in reading skills, to establish relationships between auditory cortex anatomy and individual differences in reading, to examine familial and intergenerational similarities in auditory cortex structure, and to examine intergenerational effects of parental brain structure on children’s reading. Our main findings are summarized in Figure 8, and as follows.

**Figure 8.**
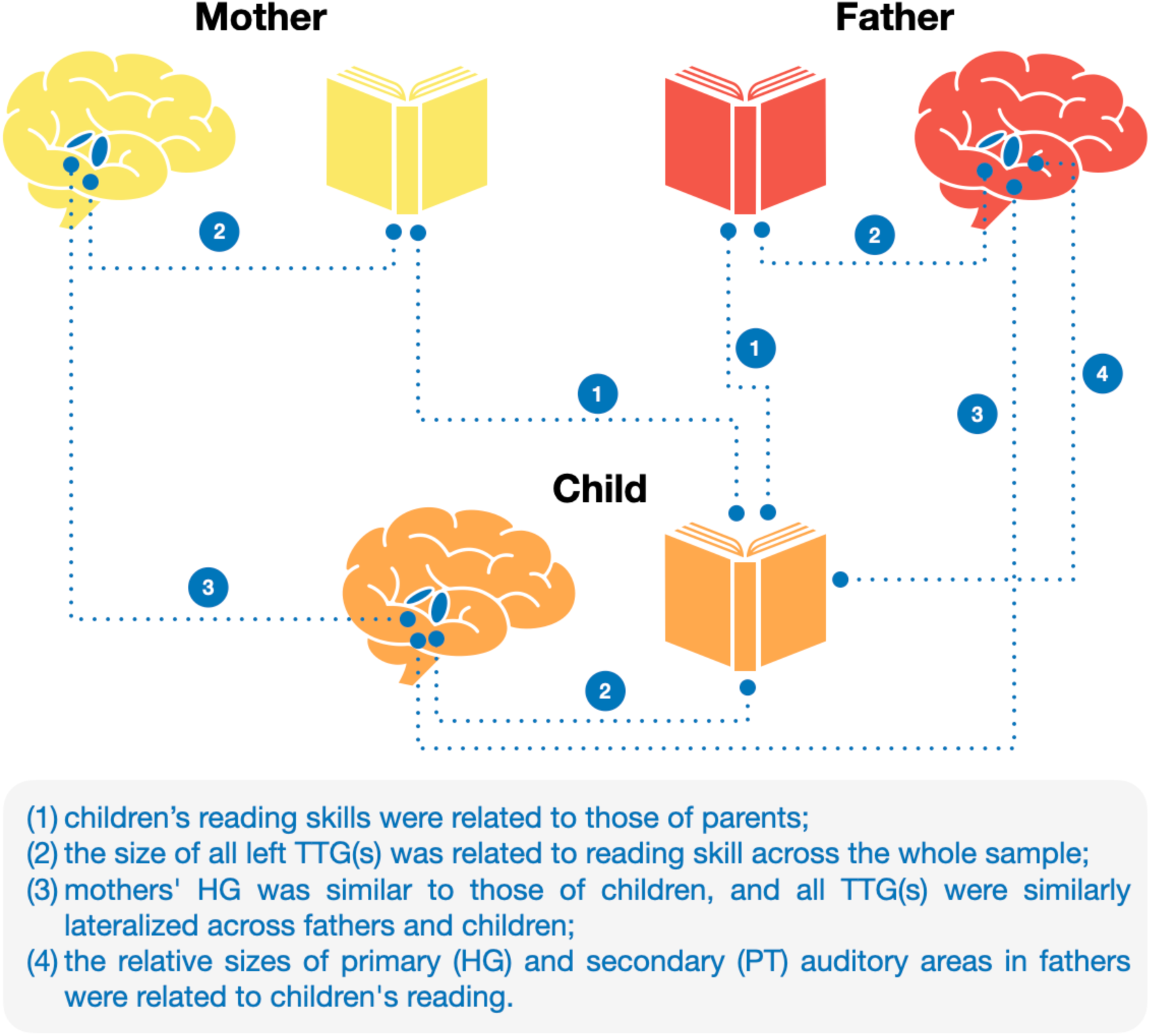
A schematic representation of the most important findings, as discussed in the text.

First, (1) we found that children’s reading skills were significantly related to those of parents, more so for mother-child pairs than for father-child pairs. Interestingly, they were observed for reading subtests which target phonological decoding skills rather than lexico-semantic, or sight-word reading. This finding is in line with twin studies showing higher heritability of phonological aspect of reading rather than lexically-mediated ones (Gay and Olson, 2003), and a recent genome-wide association study in which a higher proportion of non-word reading variability was accounted for in comparison to word-reading (Eising et al., 2022).

Furthermore, given that most of the mothers participating in the study were homemakers (21 out of 33, see Table S1), this finding might also reflect the amount of time the mothers spent with their children, and thus suggest an environmental (rather than genetic) transmission of reading skill. Alternatively, it could reflect genetically-driven parent-of-origin effects, or genomic imprinting (Lawson et al., 2013), which we discuss in more detail below (see Section 4.3).

Next, using newly developed toolboxes (Dalboni da Rocha et al., 2023, 2020), we performed detailed segmentations of gyri in the superior temporal plane, i.e., the Heschl’s gyrus and further TTG(s), when present, and found that:

(2) across the whole sample, volume and surface area of all TTG(s) in the left hemisphere correlated with individual differences in speeded phonemic decoding;
(3) there were structural brain similarities for parent-child pairs in the 1^st^ TTG (Heschl’s gyrus, HG), and in the lateralization of all TTG(s) for father-child pairs;
(4) the relative sizes of HG (including primary) and PT (consisting exclusively or mostly of secondary) auditory areas in fathers were associated with offspring reading ability.

Each of these points is discussed in more detail below, in Sections 4.1 through 4.4:

### 4.1 TTG(s) and reading abilities

Across our whole sample, individual differences in performance on speeded phonemic decoding (T-PDE) (Torgesen et al., 1999) was related to the structure of the auditory cortex when a stricter statistical correction was applied; three other tests, of speeded sight-word decoding (T-SWE) (Torgesen et al., 1999), untimed sight-word decoding (WRMT-WID) (Woodcock, 1998), and untimed phonemic decoding (WRMT-WA) (Woodcock, 1998), showed relationships with the same neuroanatomical measures when uncorrected statistics were used. Thus, we observed that both word (WRMT-WID, T-SWE) and pseudo-word reading (T-PDE, WRMT-WA) were related to some degree to TTG(s) anatomy. However, the task that assessed how well participants could pronounce syllables and pseudo-words and use their phonetic decoding skills and knowledge of grapheme–phoneme correspondence (T-PDE), was more strongly related to TTG(s) anatomy (see Figure 3B). This relationship was stronger for children than for adults, possibly pointing to developmental changes of the nature of reading throughout development: the reliance on auditorily-mediated processes is stronger in beginning stages of reading (i.e., during early school years).

This interpretation is consistent with longitudinal evidence from transparent orthographies, such as Italian, which highlight the causal importance of auditory-phonological discrimination for the development of phonological decoding skills. For example, Bertoni and colleagues (2019) showed that preschoolers’ ability to auditorily discriminate linguistically similar speech sounds predicted later phonological decoding speed, with future good and poor readers differing in their speech-sound discrimination abilities as early as kindergarten. These findings suggest that the link we observe between TTG(s) anatomy and phonological decoding might generalize across orthographies. Moreover, in a related line of research Tallal and colleagues (1980), showed that deficits in rapid auditory temporal processing could impair the development of phoneme and syllable perception, a key precursor of reading. This finding was further supported by Gaab et al. (2007), where dyslexic children exhibited altered neural activation during rapid acoustic processing which could be improved with targeted remediation. Moreover, Goswami’s temporal sampling framework (Goswami, 2011) emphasizes the role of rhythmic and temporal sampling of auditory input in dyslexia, and research by Ghesquière and colleagues (e.g., Van Ingelghem et al., 2001) demonstrates that auditory temporal processing impairments are common in dyslexia and related to phonological development. Together, these findings support the idea that structural variability in auditory cortex, specifically TTG morphology, may index individual differences in neural systems underpinning temporal and phonological processing critical for reading acquisition.

Notably, rapid automatized naming (RAN) skills, as measured by the RN-LTR task (Wolf and Denckla, 2005), showed no relationships with any neuroanatomical measures of the TTG(s), even at the uncorrected threshold. This dissociation between TTG(s) anatomical correlates of phonological processing skills *versus* RAN skills is consistent with the double-deficit hypothesis of dyslexia (Wolf and Bowers, 1999). According to this hypothesis, reading difficulties may be caused by impairments in either naming speed or phonological processing, with individuals with a “double-deficit” having more reading problems than those with a single deficit. Neuroanatomical evidence for such a dissociation has been provided by both structural (Leonard et al., 2006) and functional (Norton et al., 2014) data (although Norton and colleagues did not observe the functionality of TTG(s) to be related to the degree of phonological impairment), and our data provide further evidence for the dissociation.

In terms of the precise characteristics of the TTG(s) that were related to the variability in phonological skills in our sample, we observed positive relationships between the size (volume and surface area) of all identified TTG(s) in the left hemisphere and the relevant reading tasks. Furthermore, inspection of the results without FDR correction for multiple comparisons (Figure S 1) showed that both variables describing the 1^st^ left gyrus (volume and area of the left HG), as well as the overall shape (which includes information on the number of identified gyri as indexed by the variable ‘left all TTG(s) shape’) contributed to the ‘left all TTG(s) volume’ and ‘left all TTG(s) area’ results. To reiterate, Heschl’s gyrus (1^st^ TTG) usually hosts primary auditory processing, and gyri posterior to it and belonging to planum temporale tend to underlie secondary auditory processing. We show that it is the *combined* anatomical variability of all TTGs (HG and further TTGs, when present) that are important to reading and not either of them separately: larger and more numerous gyri in the left superior temporal plane are associated with better reading skills. This finding replicates recent results showing that in pre-readers, the surface area of left HG as well as the duplication patterns of the left TTG(s) positively predicted later word reading (Blockmans et al., 2023). The finding is also consistent with higher incidence of duplicated TTGs in relation to individual differences in non-native speech sound learning (Golestani et al., 2007), and in expert phoneticians (Golestani et al., 2011), i.e., it extends those results to reading, at the same time underscoring the importance of phonetic processing in reading. Notably, in the present results, the relationship between TTG(s) anatomy with reading was stronger than that of PT anatomy (which includes additional TTGs) with reading, indicating the specific importance of gyri within the PT for skilled reading. This is somewhat in contrast to increased activations in the PT being impacted by reading acquisition (Dehaene et al., 2015). However, as discussed elsewhere (Kepinska et al., 2025), the properties of gyri relative to sulci, such as greater neuronal density, cytoarchitectural differences or connectivity properties, may make them more likely to be associated with better auditory or higher-level linguistic skills than neighboring sulcal cortex, and therefore functional data depending on template registration and averaging of individual anatomy may miss activity patters specific to gyri *within* the PT.

Of note, none of the neuroanatomical variables describing the lateralization of TTG(s) anatomy were significantly associated with the reading measures collected, nor did the indices of the right hemisphere gyri survive correction for multiple comparisons. While a negative relationship between the shape of the right TTG(s) and reading ability might have been expected based on the results of Altarelli et al. (2014) and Serrallach et al. (2016), such an association was not present in our data. This may be due to the limited number of (very) poor readers in our sample, and as such, does not contradict the previous findings: a positive association with the overall size and gyrification of the TTG(s) in the left hemisphere and reading in our sample of predominantly typical readers may be a ‘flip side’ of the right hemisphere correlates of impaired reading.

### 4.2 Familial similarity of the TTG

Regarding TTG(s) similarity between family members, we observed intergenerational effects in HG and in the lateralization of all identified TTG(s). Intergenerational similarity between parents and children in measures of volume, surface area, and thickness were restricted to right HG, whereas shape was only significant in left HG. Both the HG and TTG(s) lateralization findings were confirmed to be significantly more likely for parent-child pairs than for random adult-child pairs, confirming that the observed effects were not due to general similarities of the brain regions studied across individuals.

#### 4.2.1 Right HG – volume, area and thickness

The structure of the right HG was similar in terms of volume, area, and thickness between mothers and children, and only in terms of thickness between fathers and children. HG houses the primary auditory cortex, but its size and shape have also been shown to be related to higher-level cognitive functions, such as musicianship (Schneider et al., 2002) and phonetic learning skill and expertise (Golestani et al., 2011, 2007). Research has suggested that the right HG uniquely underlies processing of pitch direction (Johnsrude et al., 2000), and that its larger volume is an anatomical marker of absolute pitch (Wengenroth et al., 2014). The thickness of the right HG has been specifically associated with the severity of auditory verbal hallucinations in schizophrenia patients (Chen et al., 2015). At the same time, data from our laboratory suggest that right HG surface area is positively related to individual differences in foreign language aptitude (Ramoser et al. *under review*). In the context of reading, Ma and colleagues (2015) showed a relationship between the thickness of the right HG and of surrounding areas and dyslexia, and dyslexia risk was significantly associated with polygenic risk score of bilateral HG thickness (Gialluisi et al., 2021). Whether and how individual differences in musicality, pitch perception, and language aptitude may be mediated by parent-child concordance in HG morphology requires further research. So far, however, our data do not suggest that variability in children’s reading ability can be related to the morphology of their own right HG, despite the apparent parent-offspring similarities (see also Section 3.5 on intergenerational effects on reading skills and the discussion of these findings below).

#### 4.2.2 Shape of the left HG

We also observed that the shape of mothers’ left HG was similar to that of children’s left HG. The shape of the left (but not the right) HG has previously been related to phonetic learning ability, with good phonetic learners having a higher probability of having a duplicated TTG (Golestani et al., 2007), as well with phonetic expertise (Golestani et al., 2011). Phonetic learning might be an ability intergenerationally transmitted by mothers through mother-child concordance of the shape of the left HG. To confirm this, future research should employ phenotypic testing of auditory learning abilities in both parents and offspring. Incidentally, gyrification patterns of the HG have been related to schizophrenia symptoms and suggested to be associated with susceptibility to psychopathology (Takahashi et al., 2021). Schizophrenia is a partly genetically transmitted condition (Sullivan et al., 2003), with the mother’s disorder determining the high-risk status of the offspring (Niemi et al., 2005). Mother-child similarities in HG shape may provide novel neurobiological support for maternal transmission patterns in schizophrenia, but the status of HG shape as a potential endophenotype of schizophrenia requires further research with affected individuals.

#### 4.2.3 Lateralization effects

Leftward asymmetry of the temporal speech region has been well established in the literature ever since the work of Geschwind and Levitsky (1968) (Marie et al., 2015; Penhune et al., 1996), and has been more recently confirmed by large-scale neuroimaging efforts (Kong et al., 2018). In our data, both adults and children showed a leftward asymmetry in the volume and area of HG and TTG(s), whereas the thickness of HG and TTG(s) was symmetrically distributed across the hemispheres (see Table 1). This is partially consistent with previous asymmetry investigations of auditory cortex: while Meyer and colleagues (2014) reported leftward-lateralization in terms of volume and surface area, in line with our current findings, they found a clear rightward asymmetry for thickness of the auditory regions (see also Kong et al., 2018 for similar results). This discrepancy may be due to different segmentation approaches (atlas-based *versus* individual anatomy-based), or to exclusively investigating HG (as opposed to all TTGs). We found the degree of lateralization of the surface area of TTG(s) to be similar within families, and driven by the similarity between the TTG(s) lateralization indices of fathers and their children. Lack of (i.e. symmetrical) or even reserved asymmetries of the temporal speech region have previously been reported as male-specific neural markers of dyslexia (Altarelli et al., 2014). Our results of intergenerational similarity of TTG(s) lateralization (Section 3.4.3) suggest that these dyslexia-related individual differences in structural asymmetries may be transmitted from fathers to offspring. Genetic influences on the structural lateralization of the temporal speech regions have been proposed in the literature (e.g., Eckert et al., 2002), but despite the findings that individual differences in cerebral lateralization are highly heritable (Francks, 2015), Bishop (2013) proposes that the degree of lateralization might also be driven by environmental factors. Because intergenerational similarities reflect both genetic and environmental factors common to families, our results may reflect both.

### 4.3 Parent-of-origin effects

Most of the intergenerational effects reported here were specifically patrilineal or matrilineal (with the exception of the parent-offspring similarities in average thickness of the right HG). Their basis could be environmental in nature (see, for example, Feldman (2023) for a discussion of paternal contributions to development), or an unintended by-product of our particular sample. They could also potentially be due to genetic parent-of-origin effects, which refer to phenotypic effects of an allele depending on whether it is inherited from the mother or father, and which play an important role in the genetic architecture of complex traits (Lawson et al., 2013). Such parent-of-origin effects are common in humans (Mozaffari et al., 2019), and indeed Goos et al. (2006), based on animal studies, clinical data and intergenerational analyses of behavioral data, suggest that they are “influential in brain development, with the maternal genome playing a disproportionate role in the development of the cortex” (p. 19). Our data suggest that the structure of the auditory cortex manifests such a parent-of-origin effect to some extent, and that maternal influences may indeed be more important in the development of auditory regions than paternal influences (for a further discussion of the molecular basis of the parent-of-origin effect see e.g., Hager et al., 2008; Lawson et al., 2013).

### 4.4 Intergenerational effects on reading ability

Given that none of the intergenerational TTG(s) similarity effects reported in Section 3.4 were also found to be related to reading ability (Section 3.3), we took an exploratory approach to determining whether (and how) neuroanatomical characteristics of mothers’ or fathers’ TTG(s) could predict children’s reading. The structure of the left HG was found to be the best predictor of children’s reading for fathers’ data (no neuroanatomical markers of mothers were found to be related to children’s reading). Surprisingly, fathers with a statistically smaller left HG had children who were better readers. This finding is counterintuitive because, to our knowledge, no studies have reported a negative association between left HG size and reading ability. Our follow-up analyses revealed that, in fact, it was the relative size of fathers’ HG (including primary) to secondary (PT) auditory areas that explained most variance in children’s reading (small HG:PT ratio was related to better reading). Moreover, in boys specifically, fathers’ large PT was related to better reading. These results may be related to the sex differences in the auditory cortex often observed in the literature. As mentioned above, Altarelli and colleagues (2014) found that an increased number of right TTG duplications and altered asymmetry of the PT are associated with dyslexia but only in boys; Sutherland et al. (2012) found that the sex-specificity of the association between gray matter probability (likely reflecting volume) of left Heschl’s gyrus and auditory processing differs across development (i.e., is only apparent during early adolescence).

The direction of our HG:PT ratio finding does not align with that observed in developmental disorders reported by Serrallach and colleagues (2016): diminished HG:PT ratios were reported for adults with ADHD and ADD (2022), and for children with ADHD, ADD and – crucially – with dyslexia. One could expect that an association between parental brain structure and children’s reading should follow the same direction (i.e., that a small HG:PT ratio should be related to worse reading outcomes), but our data show the opposite pattern. This might have to do with the specific, indirect nature of intergenerational effects investigated here. We related children’s behavior to parents’ brain measures, and these associations might show different patters due to, for example, neuroplasticity mechanisms operating throughout the lifespan. The inconsistent direction of our and of previous results may also arise from a non-linear relationship between reading level and HG:PT ratio; it could for example be that there is an inversed u-shaped function between HG:PT surface area (of fathers) and reading level. In turn, this could arise from the fact that intergenerational anatomical underpinnings of dyslexia may be different than those underlying healthy variability in reading skill (in other words, similar anatomical features may exist, but these regions may function differently, or display in functional and/or structural connectivity). Our finding of a father-specific association between parental left auditory cortex anatomy and offspring reading ability may, furthermore, add to the body of knowledge showing differential patterns of association between auditory cortex anatomy and cognitive abilities across sexes. However, the exact mechanisms by which paternal HG and PT are related to offspring reading, and whether there may be third variables that explain this relationship, should be explored in future research.

### 4.5 Precision imaging as a window into the neurobiological foundations of cognition and its intergenerational transmission

The present study contributes to a growing movement toward “precision imaging” approaches in cognitive neuroscience, which emphasize detailed, anatomically grounded analyses of individual differences in brain structure. Rather than relying solely on automatically extracted whole-brain estimates such as cortical thickness or surface area, measures that typically show small effect sizes and require very large samples (Marek et al., 2022), our study focused on a region of high anatomical variability and functional relevance for reading: the transverse temporal gyrus (TTG). Recent evidence suggests that macro-anatomical features such as the shape of certain gyri or their configuration may more directly reflect developmental mechanisms and functional specialization (Bouhali et al., 2024; Cachia et al., 2021; Gratton et al., 2022), and may therefore offer a more powerful window into brain-behavior associations in smaller, well-characterized samples. In line with this, our multimodal approach incorporated complementary measures: volume, surface area, cortical thickness, and shape of the TTG(s), each indexing distinct biological underpinnings. For example, surface area is considered more strongly heritable and shaped by early developmental processes, whereas cortical thickness may reflect later maturational or experience-dependent changes (Grasby et al., 2020). Shape descriptors, such as the number and configuration of TTGs, capture qualitative anatomical variation not reducible to size, and may provide particularly rich insights into familial similarity and cognitive outcomes. While most of the current precision imaging methods often require manual segmentation and modest sample sizes, our study shows that automated segmentation tailored to specific regions (here: the auditory cortex) could bridge high-resolution phenotyping with large-scale population studies. Integrating these approaches represents a promising path forward for understanding the neurobiological foundations of cognition and its intergenerational transmission.

### 4.6 Limitations and conclusions

Given the limited prior research on parent-offspring relationships with regards to reading and neuroanatomy, the precise mechanisms underlying our findings remain uncertain; however, it is reasonable to hypothesize that a convergence of genetic, prenatal, and postnatal environmental factors contribute to the observed effects. Importantly, the findings reported here are correlational in nature. While our approach integrates detailed neuroanatomical and behavioral data and employs rigorous statistical analyses, the cross-sectional and observational study design does not permit any causal conclusions. That is, we cannot determine whether parental brain structure *causally* influences children’s brain anatomy or reading abilities, or whether observed relationships reflect shared genetic or environmental factors, or a combination of both. Furthermore, given that our study was limited to genetically-related families, future work will benefit from including *in vitro* fertilization and adoptive families to isolate distinct genetic, prenatal, and postnatal environmental influences on auditory cortex structure and related (reading) phenotypes. In addition, the age of the sample studied here most participants being already readers, either for a few years (children) or for many years (parents), further limits our ability to determine the directionality of the relationship between auditory cortical neuroanatomy and phonological reading skills. While our results reveal associations, they do not clarify whether structural features influence reading development or are shaped by it. However, prior studies in pre-reading children (Blockmans et al., 2023; Clark et al., 2014; Kuhl et al., 2020) have suggested that specific aspects of auditory cortical anatomy may *precede and predict* future reading abilities, supporting a potentially causal role. Similarly, emerging evidence from neuromodulation studies provides experimental support for a causal role of the auditory cortex in reading-related skills (see Turker and Hartwigsen, 2022, 2021 for overviews). Nevertheless, anatomical variation alone is likely not deterministic. Recent intervention work (Bertoni et al., 2024) shows, for example, that just a few hours of multisensory video-game training can markedly improve phonemic perception in at-risk pre-readers, suggesting that behavioral plasticity can override anatomical predispositions. Further, as our sample was relatively homogeneous and skewed toward higher SES, with many parents having received tertiary or postgraduate education, future research should also strive to include more socioeconomically diverse populations. Finally, the relatively small sample size and exploratory nature of the study underscore the need for further research with a larger pool of participants. Nevertheless, we provide novel insights into the neural underpinnings of reading ability in the auditory cortex, children’s skills in relation to parental reading, and neuroanatomy.

## 5 Acknowledgments

Data collection was funded by the National Institutes of Health (NIH) under grant K23HD054720 to FH. The authors gratefully acknowledge support by the NCCR Evolving Language, Swiss National Science Foundation (SNSF) Agreement #51NF40_180888, and by the SNSF grant #100014_18238. OK was funded by the Marie Jahoda Stipendium from University of Vienna. FB was successively supported by NIH R01HD094834, the Institute of Convergence ILCB (supported by grants from France 2030 ANR-16-CONV-0002 and the Excellence Initiative of Aix-Marseille University A*MIDEX), and the French Foundation for Medical Research (*Fondation Médicale pour la Recherche*, FRM ARF202209015734).

## 7 Supplementary Materials

### 7.1 Participants

**Table S1.**
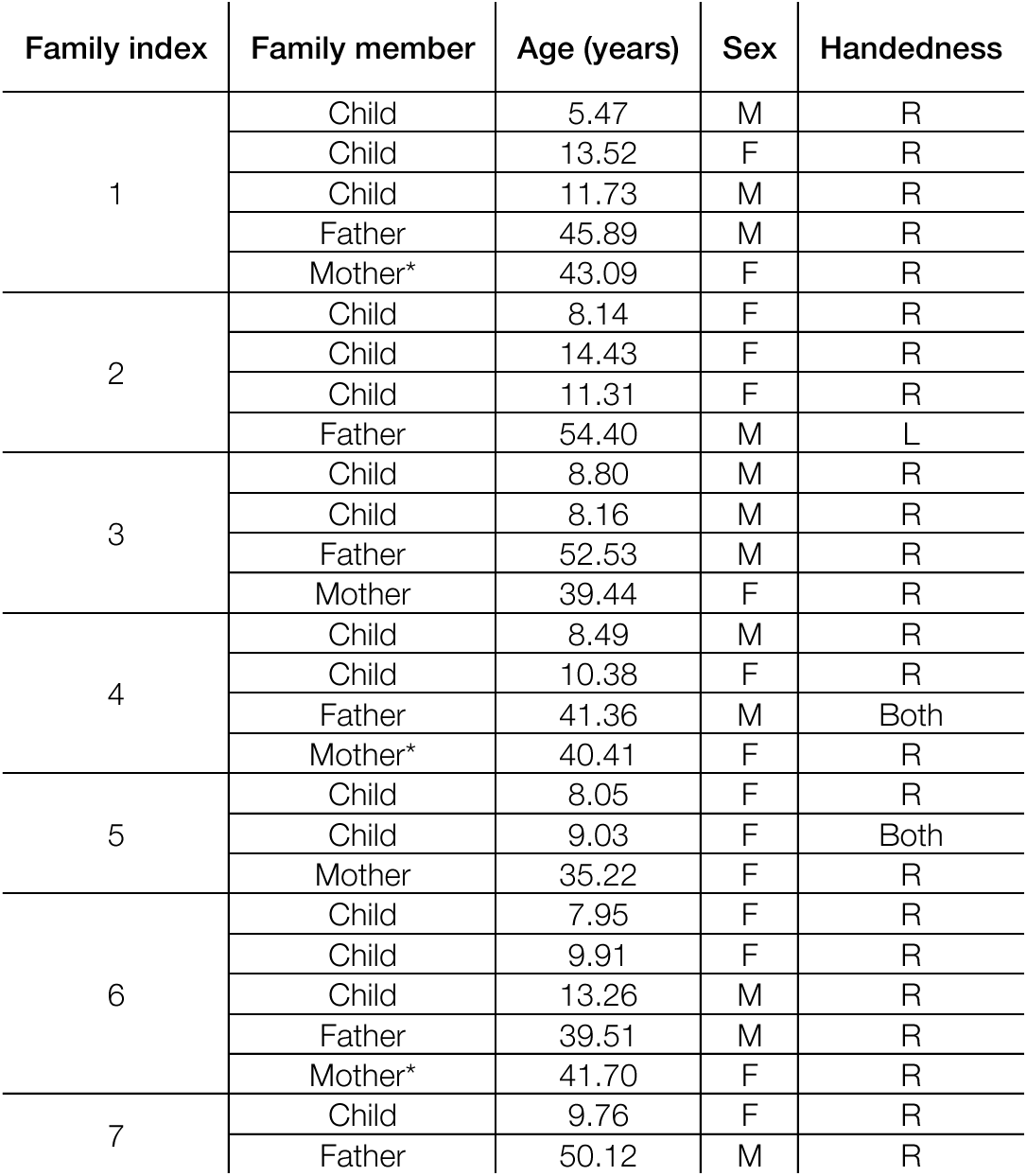
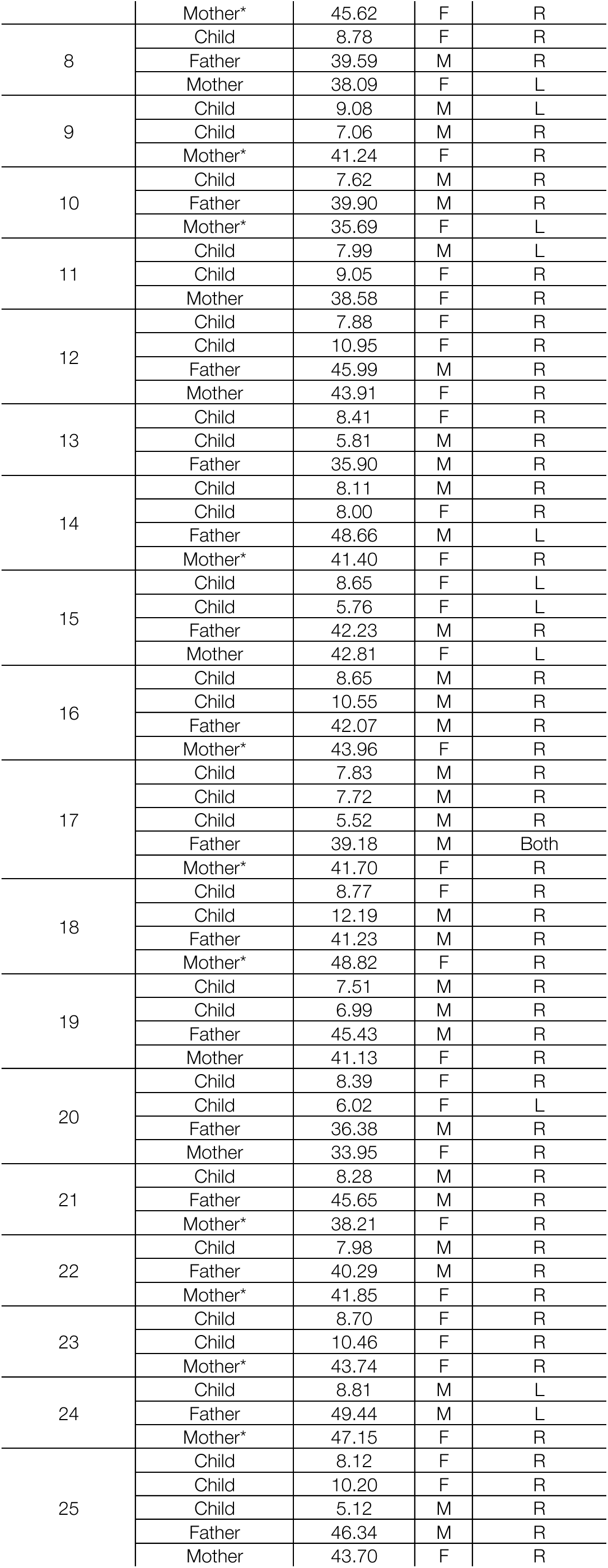
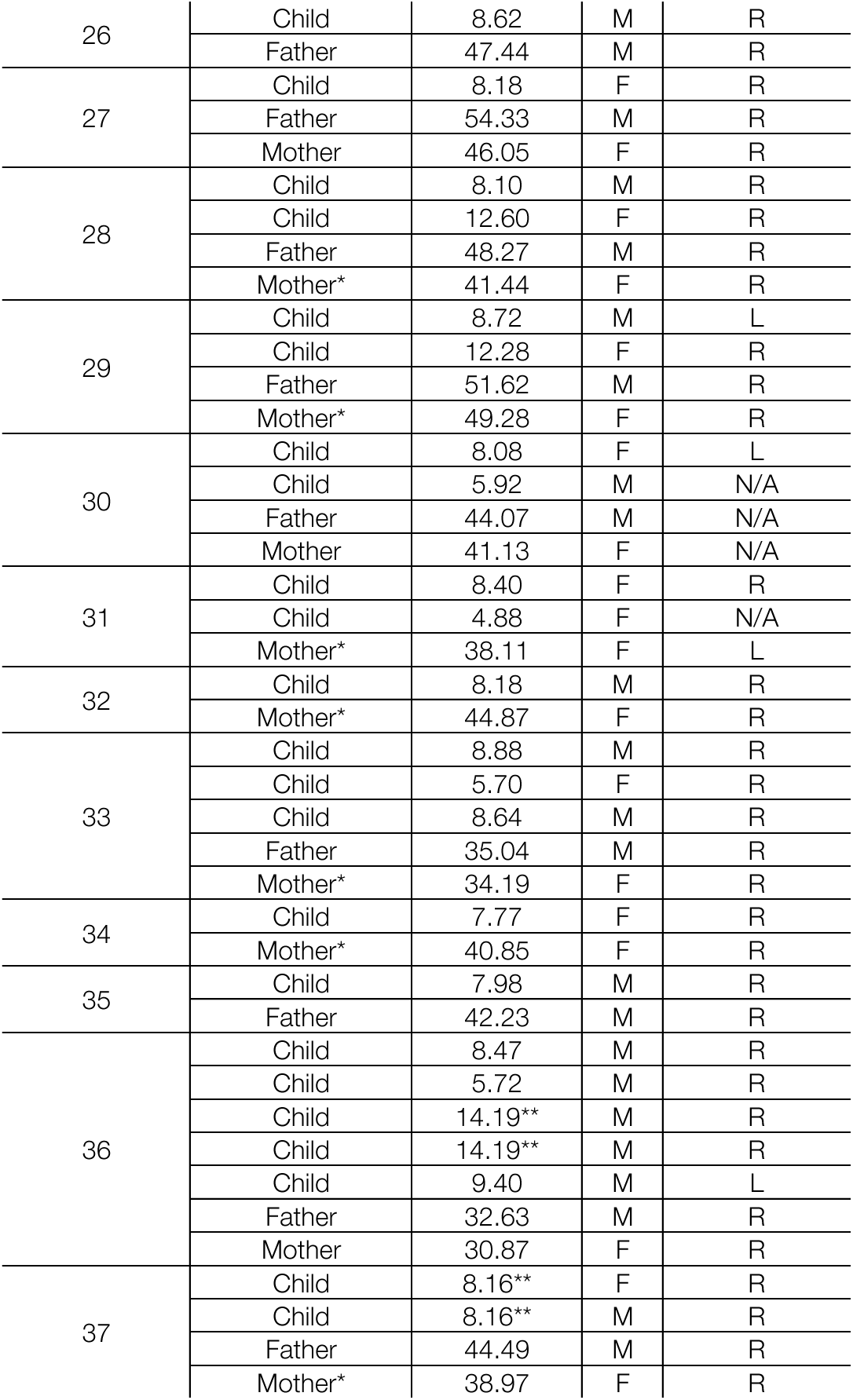
Participant characteristics including family ID, age, sex and handedness. (*) denotes mothers who declared to be homemakers; no father declared to be homemaker. (**) denotes twins.

### 7.2 TTG and reading measures (uncorrected)

**Figure S1.**
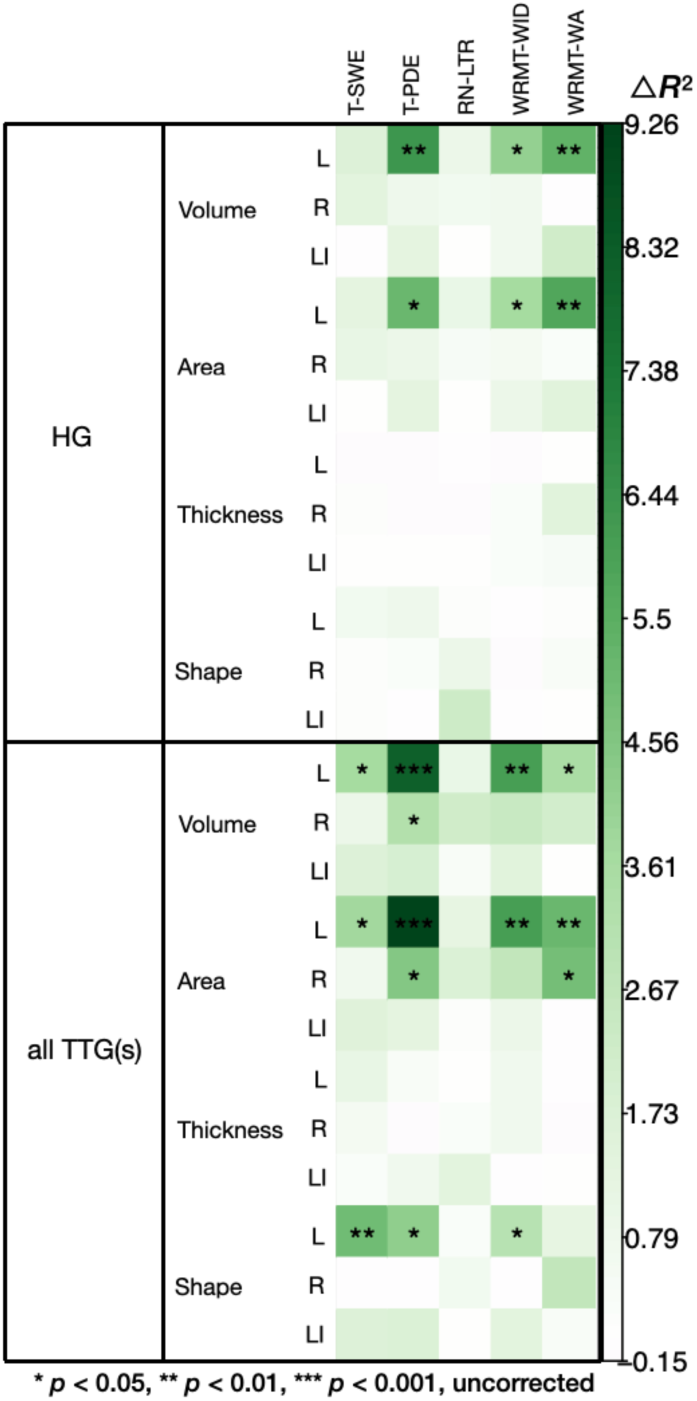
Relationship between TTG anatomy and the performance on five reading tests (T-SWE, T-PDE, RN-LTR, WRMT-WID and WRMT-WA) across the whole sample obtained from comparing models with demographic variables to models including neuroimaging data. The intensity of the color represents increase in *R*^2^ values between a model with demographic variables only and a model additionally including mothers’ or fathers’ corresponding test scores; *p*-values were obtained from likelihood ratio tests used to compare the models and are not corrected for multiple comparisons (compare with Figure 3B, where FDR-corrected *p*-values are presented).

### 7.3 Intergenerational effects on reading ability

**Figure S2.**
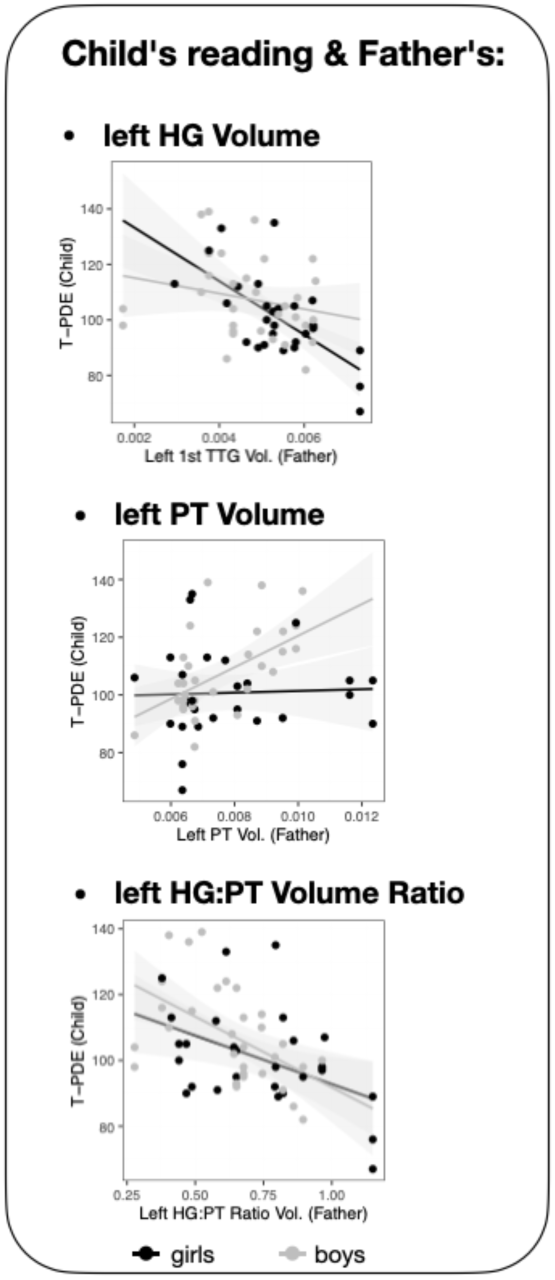
Children’s reading skills as a function of fathers’ cortical volume variables (of the left HG, left PT and left HG:PT ratio).

Similar to surface area results, fathers’ PT volume alone did not significantly predict children’s T-PDE scores (β = 0.16, *t* = 0.97, *p* = .34), but fathers’ HG:PT volumes ratio did significantly predict children’s reading (β = −0.44, *t* = −3.26, *p* = .005). Permutation analysis confirmed familial specificity of this effect (*p* = .0006). Directly comparing the models predicting children’s T-PDE with either father’s HG volume, or fathers’ HG:PT volumes ratio as independent variables showed that the model with HG:PT ratio offered a better fit to the data with △*R*^2^*_Marg._* of 2.88% (△*AIC* = −1.88). The effect was not sex specific: a nested model comparison with a likelihood ratio test did not show a better fit for a model additionally including an interaction term between fathers’ HG:PT volumes ratio and child’s sex (Χ^2^ = 1.85*, p* = 0.17).

### 7.4 Summary of all conducted analyses

| ANALYSIS | DEPENDANT MEASURE(S) | INDEPENDENT MEASURE(S) | COVARIATES OF NO-INTEREST | QUESTION | RESULTS |
| --- | --- | --- | --- | --- | --- |
| 1. Intergenerational transmission of reading skills (Section 3.2) | children's performance on five reading and reading-related tests (T-SWE, T-PDE, RN-LTR, WRMT-WID and WRMT-WA) | a. mothers' performance on the respective reading tests<br>b. fathers' performance on the respective reading tests | age<br>sex<br>SES | Is children's reading related to parents' reading? | Yes (Figure 3A):<br>a. children's performance on T-PDE, WRMT-WID and WRMT-WA was positively related to mothers' performance on the respective reading tests<br>b. children's performance on T-PDE, WRMT-WID was positively related to fathers' performance on the respective reading tests |
| 2. TTG(s) and reading measures (Section 3.3) | whole sample's performance on five reading and reading-related tests (T-SWE, T-PDE, RN-LTR, WRMT-WID and WRMT-WA) | neuroanatomical measures describing the structure of the TTG(s):<br>a. left and right HG (volume, area, thickness, shape)<br>b. left and right all TTG(s) (volume, area, thickness, shape) | age [linear and quadratic term]<br>sex<br>SES<br>handedness<br>total intracranial volume | Are reading skills related to the structure of the TTG(s)? | Yes (Figure 3B):<br>T-PDE scores were positively and significantly related to the left all TTG(s) volume ($\Delta R^2_{\text{Marg.}} = 8.15$, $p_{\text{FDR}} = .02$) and surface area ($\Delta R^2_{\text{Marg.}} = 9.26$, $p_{\text{FDR}} = .02$) |
| 2a. Follow-up analysis | whole sample's reading measures which were significantly related to 'all TTG(s)' volume and area in Analysis 2 above (T-PDE) | volume and area of the left PT: | age [linear and quadratic term]<br>sex<br>SES<br>handedness<br>total intracranial volume | Are reading skills related to the structure of the TTG(s) specifically, or rather to PT anatomy (which contain the additional TTG(s))? | Yes:<br>'all TTG(s)' measures were better at explaining reading skills than PT measures, with $\Delta R^2_{\text{Marg.}}$ of 4.39% ($\Delta\text{AIC} = -7.78$) and 3.55% ($\Delta\text{AIC} = -6.61$) for volume and area respectively |
| 3. Familial similarity in the structure of the TTG(s) (Section 3.4) | neuroanatomical measures describing the structure of the TTG(s):<br>a. left and right HG (volume, area, thickness, shape)<br>b. left and right all TTG(s) (volume, area, thickness, shape) | family index | age [linear and quadratic term]<br>sex<br>SES<br>handedness<br>total intracranial volume | Which features of the TTG(s) anatomy are similar among family members? | a. volume, area, and thickness of right HG<br>b. shape of left and right HG<br>c. lateralization of volume and surface of all TTG(s)<br>d. thickness of the left and right all TTG(s)<br>e. lateralization of HG area |
| 3a. Follow-up analysis on Result (a) in Analysis 3 (Section 3.4.1) | children's volume, area, and thickness of the 1<sup>st</sup> right TTG (HG) [normalized for their corresponding whole hemisphere measures] | a. mothers' volume, area, and thickness of the right HG [normalized for their corresponding whole hemisphere measures]<br>b. fathers' volume, area, and thickness of the right HG [normalized for their corresponding whole hemisphere measures] | children's:<br>age [linear and quadratic term]<br>sex<br>SES<br>handedness | Is there intergenerational transmission of volume, area, and thickness of right HG? | Yes (Table 2, Figure 4):<br>a. children's volume, area, and thickness of right HG was positively and significantly related to mothers' right HG volume, area, and thickness respectively (volume: $\beta = 0.38$, $t = 2.84$, $p = .009$; area: $\beta = 0.30$, $t = 2.33$, $p = .03$; thickness: $\beta = 0.38$, $t = 2.87$, $p = .008$),<br>b. children's right HG thickness was positively and significantly related to fathers' thickness ($\beta = 0.28$, $t = 2.05$, $p = .05$), but not to other measures |
| 3b. Follow-up analysis on Result (b) in Analysis 3 (Section 3.4.2) | children's left and right HG shape | a. mothers' left and right HG shape<br>b. fathers' left and right HG shape | children's:<br>age [linear and quadratic term]<br>sex<br>SES<br>handedness | Is there intergenerational transmission of left and right HG shape? | Yes (Table 3, Figure 5):<br>a. children's left HG shape was only significantly and positively related to mothers' left HG shape ($\beta = 0.31$, $t = 2.12$, $p = .04$)<br>b. relationships between children's and fathers' HG shape were not significant<sup>2</sup> |
| 3c. Follow-up analysis on Result (c) in Analysis 3 (Section 3.4.3) | children's lateralization index (LI) of volume and area of all TTG(s) | a. mothers' LI of volume and area of all TTG(s)<br>b. fathers' LI of volume and area of all TTG(s) | children's:<br>age [linear and quadratic term]<br>sex | Is there intergenerational transmission of lateralization of all TTG(s)? | Yes (Table 4, Figure 6):<br>a. relationships between children's and mothers' all TTG LI were not significant |

<sup>2</sup> Non-significant results in gray; significant in black.

| ANALYSIS | DEPENDANT MEASURE(S) | INDEPENDENT MEASURE(S) | COVARIATES OF NO-INTEREST | QUESTION | RESULTS |
| --- | --- | --- | --- | --- | --- |
| | | | • SES<br>• handedness | | b. LI of surface area of children's all TTG(s) was positively and significantly related to fathers' all TTG(s) LI of the surface area ($\beta = 0.42$, $t = 3.00$, $p = .005$) |
| 3d. Follow-up analysis on Result (d) in Analysis 3 (Section 3.4.4) | children's thickness of the left and right all TTG(s) | a. mothers' thickness of the left and right all TTG(s)<br>b. fathers' thickness of the left and right all TTG(s) | children's:<br>• age [linear and quadratic term]<br>• sex<br>• SES<br>• handedness | Is there intergenerational transmission of thickness of the left and right all TTG(s)? | No:<br>a. mothers' measures were not significantly related to those of the children<br>b. fathers' measures were not significantly related to those of the children |
| 3e. Follow-up analysis on Result (e) in Analysis 3 (Section 3.4.4) | children's lateralization of HG area | a. mothers' lateralization of HG area<br>b. fathers' lateralization of HG area | children's:<br>• age [linear and quadratic term]<br>• sex<br>• SES<br>• handedness | Is there intergenerational transmission of lateralization of HG area? | No:<br>a. mothers' measures were not significantly related to those of the children<br>b. fathers' measures were not significantly related to those of the children |
| 4. Intergenerational effects on reading ability (Section 3.5) | children's performance on T-PDE | a. mothers' neuroanatomical measures describing the structure of the TTG(s):<br>- left and right HG (volume, area, thickness [normalized for their corresponding whole hemisphere measures], shape)<br>- left and right all TTG(s) (volume, area, thickness [normalized for their corresponding whole hemisphere measures], shape)<br>b. fathers' neuroanatomical measures describing the structure of the TTG(s):<br>- left and right HG (volume, area, thickness [normalized for their corresponding whole hemisphere measures], shape)<br>- left and right all TTG(s) (volume, area, thickness [normalized for their corresponding whole hemisphere measures], shape) | - | Is children's reading related to parents' neuroanatomy? | Yes (Figure 7A):<br>a. model with mothers' variables did not point to any features of their TTG(s) to be relevant predictors of children's reading ability<br>b. fathers' volume and area of the left HG and HG lateralization indices of volume and area had conditional permutation importance significantly different from zero |
| 4a. Follow-up analysis on Results in Analysis 4 | children's performance on T-PDE | a. fathers' left HG volume and area [normalized for their corresponding whole hemisphere measures]<br>b. fathers' lateralization of volume and area of HG | children's:<br>• age<br>• sex<br>• SES | Do the above effects survive in linear mixed effects models? What is their direction? | a. fathers' left HG volume and area were both negatively related to children's reading (volume: $\beta = -0.40$, $t = -2.91$, $p = .008$; area: $\beta = -0.46$, $t = -3.45$, $p = .002$) (Figure 7B)<br>b. fathers' lateralization of HG (both for volume and area) was not significantly related to the children's reading |
| 4b. Follow-up analysis on Results in Analysis 4a | fathers' left HG volume and area | a. fathers' left all TTG(s) shape<br>b. fathers' left PT volume and area | - | Is fathers' small left HG related to more additional TTGs, or with a larger PT? | a. No: fathers' small left HG is not related to more additional TTGs<br>b. Yes: fathers' small left HG area is related to larger PT area ($\beta = 0.18$, $t = 2.036$, $p = .05$), but volume of fathers' HG was not related to PT volume (Figure 7B) |
| 4c. Follow-up analysis on Results in Analysis 4b | children's performance on T-PDE | a. fathers' left HG area and volume<br>b. fathers' left PT area and volume<br>c. fathers' left HG:PT ratio of area and volume | children's:<br>• age<br>• sex<br>• SES | Which characteristics of fathers' auditory cortex (HG, PT or HG:PT ratio) explain most variance in children's reading? | fathers' left HG:PT ratio of area was the best predictor of children's reading (Figure 7B) |

## Footnotes

1 Note that normative data of the TOWRE and RAN/RAS subtests are only available for ages 6 to 24, and 5 to 18, respectively. Therefore, the standard scores for the available oldest age range reference were used for adult participants.

